# Cadherin Clusters Stabilized by a Combination of Specific and Nonspecific Cis-Interactions

**DOI:** 10.1101/2020.05.22.110015

**Authors:** Connor J. Thompson, Zhaoqian Su, Vinh H. Vu, Yinghao Wu, Deborah E. Leckband, Daniel K. Schwartz

## Abstract

We demonstrate a combined experimental and computational approach for the quantitative characterization of lateral interactions between membrane-associated proteins. In particular, weak, lateral (cis) interactions between E-cadherin extracellular domains tethered to supported lipid bilayers, were studied using a combination of dynamic single-molecule Förster Resonance Energy Transfer (FRET) and kinetic Monte Carlo (kMC) simulations. Cadherins are intercellular adhesion proteins that assemble into clusters at cell-cell contacts through cis- and trans- (adhesive) interactions. A detailed and quantitative understanding of cis-clustering has been hindered by a lack of experimental approaches capable of detecting and quantifying lateral interactions between proteins on membranes. Here single-molecule intermolecular FRET measurements of wild-type E-cadherin and cis-interaction mutants combined with simulations demonstrate that both nonspecific and specific cis-interactions contribute to lateral clustering on lipid bilayers. Moreover, the intermolecular binding and dissociation rate constants are quantitatively and independently determined, demonstrating an approach that is generalizable for other interacting proteins.

## Introduction

The quantitative characterization of protein interactions on membranes and at buried interfaces, including the measurement of binding constants, is a major challenge due to the limited experimental approaches capable of interrogating molecular interactions in these environments. While it is common to study interactions between extracellular regions of membrane proteins in solution, such experiments are imperfect proxies for measuring actual membrane protein interactions. Apart from the potential impact of domain isolation on protein folding and function, functionally important protein interactions and oligomerization may arise specifically due to constraints imposed by two- or three-dimensional confinement (Różycki, Lipowsky, & Weikl, 2010; Weikl, Asfaw, Krobath, Różycki, & Lipowsky, 2009). Notably, the immunological synapse is characterized by the spatial and temporal organization of proteins in the gaps between the surface of an antigen presenting cell and a T-cell (Grakoui et al., 1999; Monks, Freiberg, Kupfer, Sciaky, & Kupfer, 1998). This organization is attributed in part to the steric segregation of proteins of different sizes and to cytoskeletal interactions (Qi, Groves, & Chakraborty, 2001; Schmid et al., 2016); the understanding of the role of lateral protein interactions in this protein assembly remains incomplete. In addition to cadherins, nectins represent another class of membrane proteins whose lateral clusters mediate cell-cell adhesion (Rikitake, Mandai, & Takai, 2012). Distinct lateral (cis) and trans- (adhesive) interactions between the four members of the nectin family are associated with differentiation and tissue organization. Although it is possible to quantify trans- (adhesive) interactions (Chesla, Selvaraj, & Zhu, 1998; Chien et al., 2008; J. Wu et al., 2008), measurements of lateral interactions underlying protein clustering have been inaccessible.

In this context, cadherins pose a particular challenge. Cadherins are transmembrane proteins that mediate cell-to-cell adhesion in all tissues and regulate a range of biological processes, such as tissue rearrangement and formation, cell motility, proliferation, and signaling (Gumbiner, 1996, 2005; Niessen, Leckband, & Yap, 2011; Pla et al., 2001; Takeichi, 1995). Cadherins mediate inter-cellular adhesion by binding other cadherins on an adjacent cell surface. Notably, cadherins assemble into dense clusters at these adhesive sites, which are important for regulating the permeability of barrier tissues such as the intestinal epithelium (Brieher, Yap, & Gumbiner, 1996; O. J. Harrison et al., 2011; Y. Wu, Kanchanawong, & Zaidel-Bar, 2015). The molecular basis underlying cadherin cluster assembly is therefore of great interest because of its importance for tissue functions.

Experimental evidence supported the postulate that cadherin-mediated adhesion and clustering involves both cis- and trans-interactions between cadherin molecules on cell surfaces, where cis-interactions occur laterally between cadherin molecules on the same cell surface, and trans-interactions occur between cadherins on opposing cell membranes (Brieher et al., 1996; O. J. Harrison et al., 2011; Y. Wu et al., 2015). Early comparisons of cadherin extracellular domain adhesive activity suggested that the protein functions as a cis-dimer, and crystal structures suggested a plausible cis-binding interface (Brieher et al., 1996; O. J. Harrison et al., 2011). Moreover, mutating one or two key amino acids in the postulated cadherin cis-binding interface results in impaired intercellular adhesion and reduced cadherin clustering at cell-cell contacts (Erami, Timpson, Yao, Zaidel-Bar, & Anderson, 2015; O. J. Harrison et al., 2011; Shashikanth, Kisting, & Leckband, 2016; Y. Wu et al., 2015). However, despite experimental evidence for the importance of cis-interactions in cell adhesion, they have been difficult to investigate directly (Brieher et al., 1996; du Roure, Buguin, Feracci, & Silberzan, 2006; O. J. Harrison et al., 2011; Hong, Troyanovsky, & Troyanovsky, 2013; Indra et al., 2018; Klingelhofer, Laur, Troyanovsky, & Troyanovsky, 2002; D. Leckband & Sivasankar, 2012; D. E. Leckband & de Rooij, 2014; Shapiro et al., 1995; R. B. Troyanovsky, Indra, Chen, Hong, & Troyanovsky, 2015; R. B. Troyanovsky, Laur, & Troyanovsky, 2007; Regina B. Troyanovsky, Sokolov, & Troyanovsky, 2003; Y. Wu et al., 2015; Alpha S. Yap, Brieher, Pruschy, & Gumbiner, 1997; A. S. Yap, Niessen, & Gumbiner, 1998; B. Zhu et al., 2003). Due to the relatively weak nature of cis-interactions, traditional solution-phase studies have failed to detect them, even at high protein concentrations (Haussinger et al., 2004; Koch, Bozic, Pertz, & Engel, 1999). Furthermore, attempts to stabilize weak cis-interactions through chemical crosslinking in solution were unsuccessful (Zhang, Sivasankar, Nelson, & Chu, 2009).

Computational models of cadherin binding subsequently suggested that the reduction of configurational and orientational entropy under two- and three-dimensional confinement could potentiate cis-interactions. Specifically, the models predicted that tethering cadherin extracellular domains to a two-dimensional (2D) surface, such as a supported lipid bilayer or cell membrane would increase the effective binding affinities of both cis-and trans-interactions (O. J. Harrison et al., 2011; Y. Wu et al., 2010; Y. Wu, Vendome, Shapiro, Ben-Shaul, & Honig, 2011). Unfortunately, measurements based on analyses of photon counting histograms were not sufficiently sensitive to detect cis-interactions between E-cad extracellular domains on supported bilayers at modest cadherin surface concentrations outside of the cell adhesion zone (Biswas et al., 2015). However, the prediction that membrane-tethered cadherins can form clusters under 2D confinement was recently confirmed indirectly via single-molecule tracking, based on measurements of the diffusion of E-cadherin extracellular domains on supported lipid bilayers, over a very large range of cadherin surface coverage (Thompson, Vu, Leckband, & Schwartz, 2019). Comparisons of wild-type and cis-mutants confirmed that a specific cis-binding interface mediated clustering in the absence of *trans* interactions. Importantly, the diffusion coefficient served as a very sensitive proxy for cis-interactions, because clusters diffuse more slowly than monomers. These findings suggested that cis-interactions between E-cad extracellular domains can result in the formation of large clusters, in the absence of trans-interactions, for cadherin surface coverage above a threshold of ∼1,100 E-cad/µm^2^ (Thompson et al., 2019). However, a quantitative understanding of cis-interaction contributions to the assembly of adhesive junctions has been hindered by the lack of approaches capable of identifying and quantifying relevant binding interactions.

Here we used intermolecular single-molecule FRET microscopy to characterize the dynamic interactions between E-cad extracellular domains tethered to mobile supported lipid bilayers, while simultaneously tracking the motion of E-cad monomers and clusters to determine their diffusion coefficients, and thereby infer their hydrodynamic diameters. By comparing the behavior of wild-type E-cad to that of a mutant that is incapable of specific cis-interactions, we identified two distinct types of lateral interactions, which we attributed to nonspecific interactions (present for both wild-type and mutant E-cad) and specific interactions (present only for wild-type E-cad). The specific interactions were significantly stronger, resulting in longer intermolecular associations and a steady-state cluster distribution with a larger characteristic cluster size. Complementary off-lattice kinetic Monte Carlo simulations were performed under conditions designed to mimic the experiments. The kinetic parameters associated with the simulations were constrained by experimental values when applicable; the remaining parameters were optimized so that the steady state cluster size distributions matched those observed experimentally. The experiments and simulations were internally consistent, with a single set of parameters for all experimental conditions. These simulation results suggested that the dissociation rate for specific cis-interactions was approximately 10x slower than for nonspecific interactions under the conditions of the experiments. Thus, while associations due to nonspecific interactions were significantly weaker than cis-interactions, they were substantial and could not be ignored. The simulations also suggested that associations due to cis-interactions were more efficient and likely to occur, than nonspecific interactions. Importantly, the methods developed and employed here can be generally applied to study the dynamics of specific and nonspecific lateral interactions between a wide range of membrane proteins.

## Results

### Nonspecific and Specific Cis-Interactions Are Present in E-cad Clusters

In order to study E-cad lateral interactions under 2D confinement, donor (Alexa 555) labeled, acceptor (Alexa 647) labeled, and unlabeled E-cad extracellular domains were simultaneously bound to a supported lipid bilayer via hexahistidine-NTA associations and imaged using a prism-based total internal reflection fluorescence (TIRF) microscope. This allowed the observation of a large number of single molecule trajectories at high or intermediate protein surface coverage. Each molecular observation within each trajectory was then classified as either a high-FRET or low-FRET efficiency state (where high-FRET corresponds to a putative cis-association) based on the donor and acceptor intensities using an algorithm described previously, allowing the identification of high-FRET and low-FRET time intervals (Chaparro Sosa et al., 2018). In order to distinguish the effects of specific cis-interactions, wild-type E-cad and mutant E-cad extracellular domain constructs were used in separate experiments; this particular mutant has previously been shown to be incapable of interacting through the cis-interface (O. J. Harrison et al., 2011; Thompson et al., 2019). Therefore, at similar surface coverage, any difference in apparent interactions between the wild-type and this mutant should primarily be due to the presence or absence of specific cis-interactions.

Three conditions were studied: high-coverage wild-type (∼1,400 E-cad/µm^2^), high-coverage mutant (∼1,300 E-cad/µm^2^), and intermediate-coverage wild-type (∼1,000 E-cad/µm^2^), where these coverage values were chosen based on previous experiments that demonstrated the onset of significant clustering at surface coverages above ∼1,100 E-cad/µm^2^ (Thompson et al., 2019). These surface coverage values are physiologically relevant as they are well below the maximum local surface coverage within cell-cell adhesions of ∼49,000 E-cad/µm^2^ (Biswas et al., 2015). A total of ∼4,000 trajectories were observed, of which ∼750 exhibited FRET events, consisting of ∼85,000 total molecular observations at each of the three experimental conditions employed. Table S1 contains the exact number of total trajectories, trajectories exhibiting FRET association, and total number of displacements for each experimental condition. To permit single molecule localization, the donor-labeled E-cad concentration was kept very low as described in the Materials and Methods section. The acceptor-labeled E-cad concentration was much larger than that of the donor, allowing the observation of a large number of FRET events and to ensure that multiple acceptors were present in clusters. Due to limitations in acceptor concentration caused by the need to avoid excess background that results from direct excitation of the acceptor, unlabeled E-cad was added to reach sufficiently high surface coverages required for cluster formation.

In some trajectories, transitions between FRET states were observed, presumably indicating association and dissociation events between donor and acceptor labeled E-cad. However, many trajectories showed no FRET-state transitions, where a trajectory began by either adsorption or diffusion into the field of view in a given state and remained in that state until the trajectory ended through desorption, diffusion out of the frame, or photobleaching. Representative trajectories illustrating these different situations are shown below in Figs. 1 A-I. For example, in trajectory 1, the donor E-cad begins in the low-FRET state and appears to be diffusing quickly, based upon the large positional fluctuations. After ∼0.33 s, a transition from low to high FRET-state indicates the association of the donor E-cad with a cluster. This FRET transition coincides with a significant decrease in the positional fluctuations, consistent with the motion of a large cluster. In contrast, representative trajectory 2 exhibits no apparent FRET-state transitions. The trajectory begins in the high-FRET state and remains in this state throughout the entire trajectory. The position fluctuations are small and the molecule remains in a small, confined, region. This behavior suggests that the donor E-cad is associated with a large cluster that contains one or more acceptor E-cad molecules. Lastly, trajectory 3 remains in the low-FRET state throughout the entire trajectory, and exhibits large positional fluctuations, consistent within an unassociated monomer of donor E-cad.

**Fig. 1.**
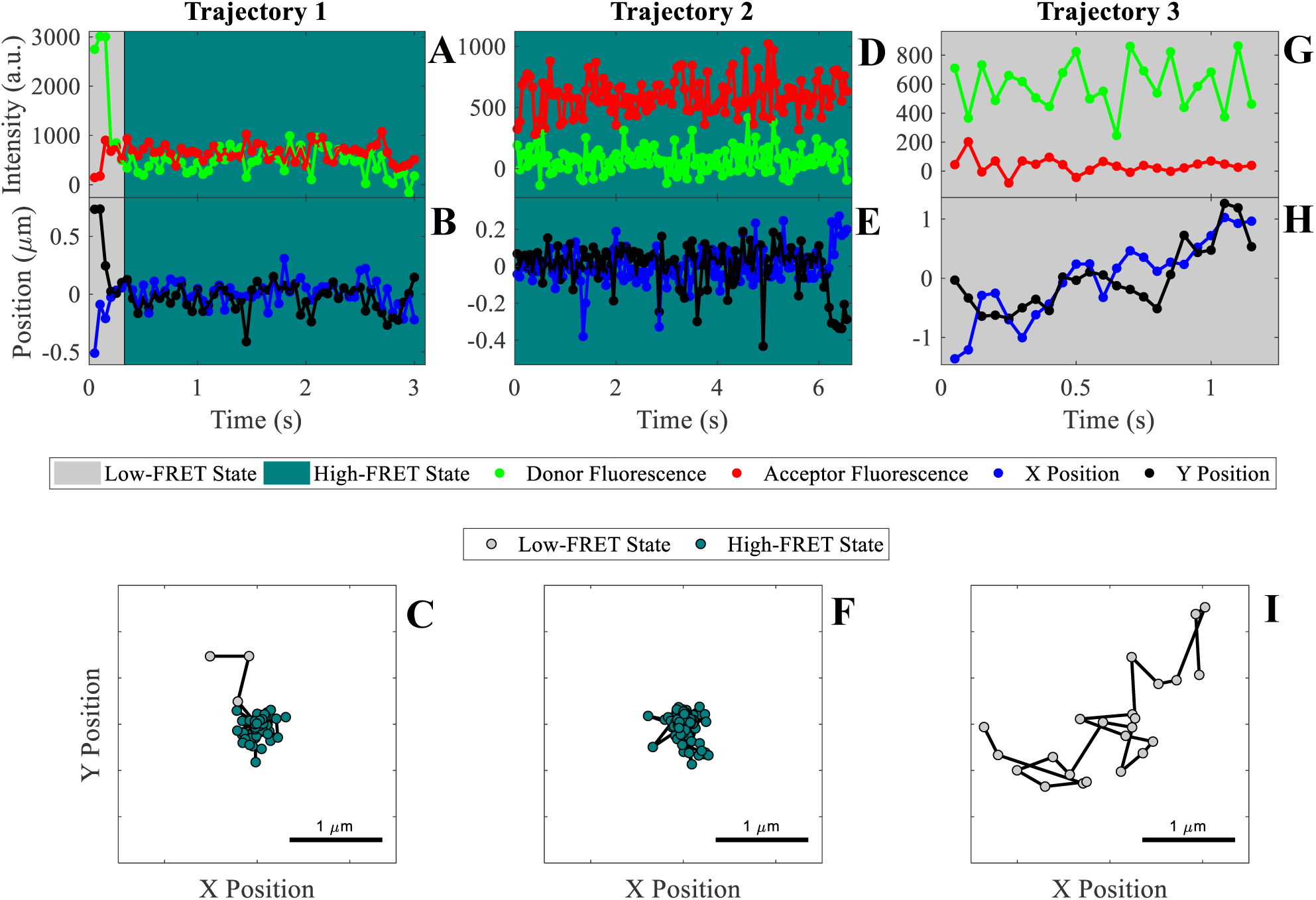
Representative trajectories for individual E-cad extracellular domains diffusing on a supported lipid bilayer. (A, D, G) Donor and acceptor intensity for the FRET pair throughout the trajectory, which is used to determine if the donor E-cad molecule is in a high-FRET or low-FRET state. (B, E, H) X and Y Cartesian coordinates for the donor or acceptor molecule over the length of the trajectory. (C, F, I) Two dimensional trajectory plots of the same trajectories, where the symbol color corresponds to the assigned FRET-state. The background of the trajectory time traces for intensity and position indicate the assigned FRET-state.

As is apparent from Fig. 1 A-I, transport properties are often coupled to the FRET-state of a molecule. This is because the FRET-state reflects the oligomeric state of an E-cad molecule, and large oligomers diffuse slower than a monomer due to increased protein-lipid interactions, which is the primary source of drag (Cai, Shashikanth, Leckband, & Schwartz, 2016). In order to assess this hypothesis and confirm that the high-FRET state does in fact correlate with protein clusters involving a donor and one or more acceptors, the average short-time diffusion coefficient 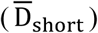 was determined for the high and low FRET-state populations independently. This was done by constructing complementary cumulative squared displacement distributions (CCSDDs) for each state, under each experimental condition, and then fitting these distributions to a Gaussian mixture model containing three terms (See Materials and Methods section for more details on distribution calculations, fitting, and 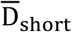 calculation). Additionally, overall CCSDDs were constructed, in order to determine overall values of 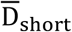 under each experimental condition. Overall CCSDDs and Gaussian mixture model fits are shown in Fig. S2. Fig. 2A-C below shows the CCSDDs for both FRET-states (at each of the three experimental conditions) with the respective Gaussian mixture model fits. Fig. 2D shows the resulting values of 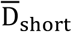 determined from the fit parameters. Table S2 shows all CCSDD fit parameters.

**Fig. 2.**
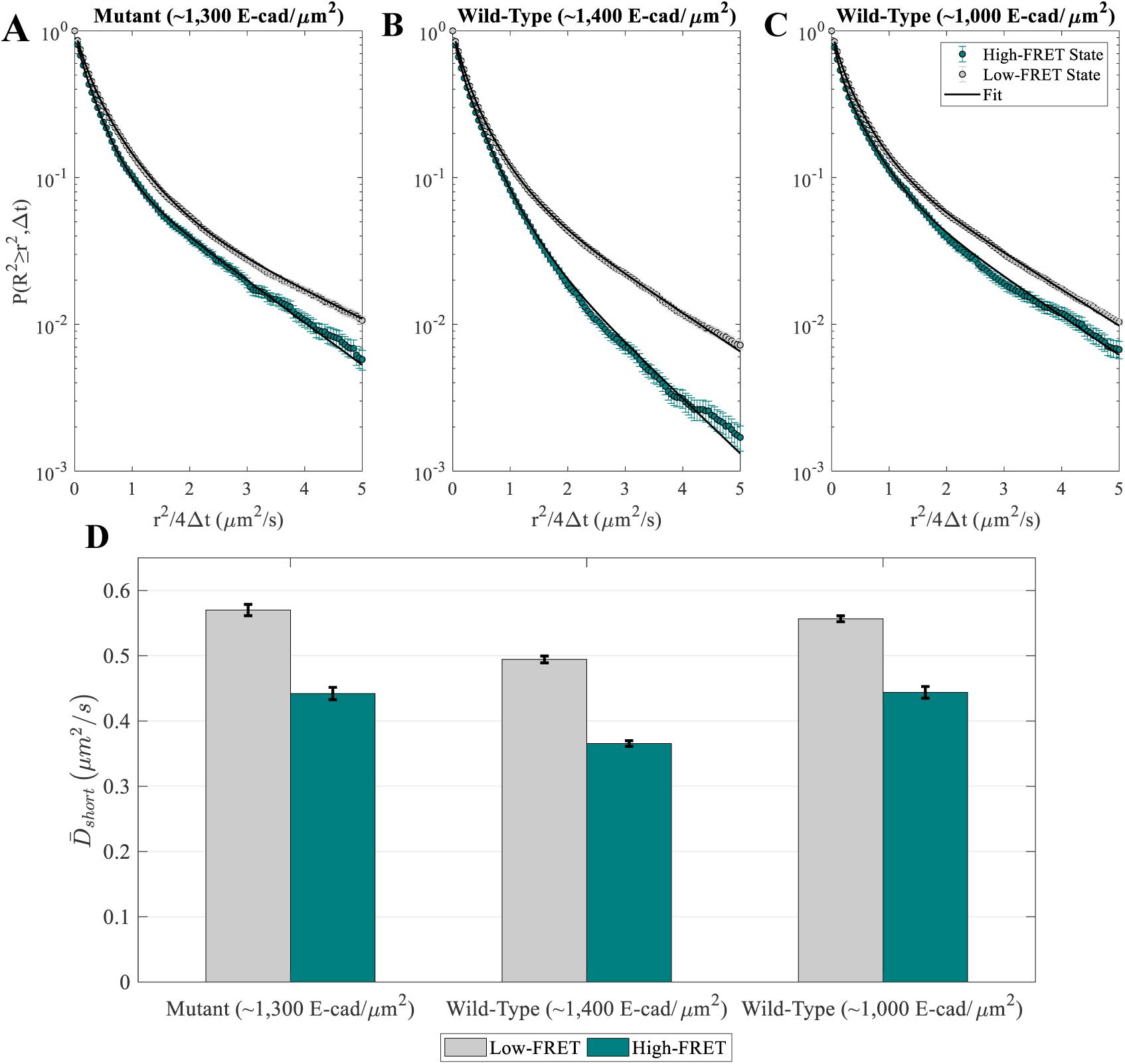
E-cad diffusion depends on FRET state and interaction capability. (A-C) Complementary cumulative squared displacement distributions in the high-FRET and low-FRET states for the mutant and two wild-type E-cad conditions, along with the respective Gaussian mixture model fits. Error bars correspond to the standard deviation of CCSDDs calculated using 100 samples using a bootstrap method with replacement. (D) 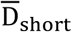 in the high-FRET and low-FRET states for the mutant and two wild-type conditions. Error bars represent the standard deviation of fitting 100 samples using a bootstrap method with replacement.

Most importantly, Fig. 2D shows that the values of 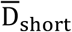 are significantly smaller for the high-FRET state relative to the low-FRET state. This behavior is consistent with the interpretation that the high-FRET state corresponds to E-cad in an associated state, where it diffuses as an oligomer or large cluster. Of course, due to the presence of unlabeled E-cad, it is possible for an E-cad donor molecule to be associated with a cluster but remain in a low-FRET state. The low-FRET state population comprises a combination of unassociated donor E-cad and donor E-cad that is associated with unlabeled E-cad; consequently, this population is more complicated to interpret. Nevertheless, the inclusion of monomers in the low-FRET state (and not the high-FRET state) is expected to result in larger values of 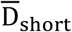 for the low-FRET state, as observed for all three experimental conditions, even for the mutant that cannot interact through the cis-interface. Importantly, the observation that the mutant also exhibits decreased diffusion in a high-FRET state (from 0.569 ± 0.008 μm^2^/s to 0.44 ± 0.01 μm^2^/s) suggests that the proteins can associate by nonspecific interactions in addition to the specific cis-binding interface expected for wild-type E-cad.

As shown in Fig. 2D, the protein diffusion associated with each of the FRET states is slowest at the higher surface coverage of wild-type E-cad. This observation is consistent with the presence of more large protein clusters than at lower surface concentrations or in the absence of specific cis-interactions. The formation of these large clusters is presumably supported by a large number of nonspecific interactions, in combination with frequent cis-interactions at the higher concentration. Interestingly, the diffusion constants of both FRET-state populations are similar for wild-type E-cad at lower surface concentration and for mutant E-cad at higher concentration. This finding suggests that the average cluster sizes are comparable in these two systems, due to a balance between the strength and frequency of the different nonspecific and specific interactions. This is consistent with previous findings that specific cis-interactions between wild-type E-cad proteins primarily affected diffusion only at surface coverages above ∼1,100 E-cad/µm^2^, while nonspecific interactions between mutant E-cad did not cause significant slowing even above this threshold (Thompson et al., 2019). Additionally, the overall 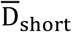 values (Table S2) show that effective total diffusion was slowest for high surface coverage wild-type E-cad, and that the overall diffusion for mutant E-cad and intermediate coverage wild-type E-cad was similar. To better understand the relationship of nonspecific and specific lateral interactions between E-cad extracellular domains, a detailed investigation of the dwell time distributions in the high and low-FRET states was performed as described below.

### Nonspecific Cis-Interactions Dissociate Faster than Specific Cis-Interactions

Classifying each observed trajectory into the high-FRET or low-FRET state provides information about the time intervals spent in each state (dwell time), in addition to the state-dependent transport properties discussed previously. The dwell times in each state contain direct information about the nature and energies of interactions. These data can be used in tandem with the transport information, which provides indirect information about clustering. Fig. 3A shows complementary cumulative dwell time distributions for the high-FRET state, under the three conditions studied (see Materials and Methods section for details on the complementary cumulative dwell time distribution calculations). For these semi-logarithmic plots, simple first-order desorption kinetics would appear as a straight line. Therefore, the curved plots presumably indicate the presence of multiple populations or modes of dissociation. Interestingly, the data obtained with the mutant E-cad at high concentration are notably less heterogeneous (i.e. less curved) than those of the wild-type E-cad at either concentration. The complementary cumulative dwell time distributions in the low-FRET state are identical for all experimental conditions, as shown as Fig. S3. This is discussed in the section concerning kMC simulations, in the context of association rate constants.

**Fig. 3.**
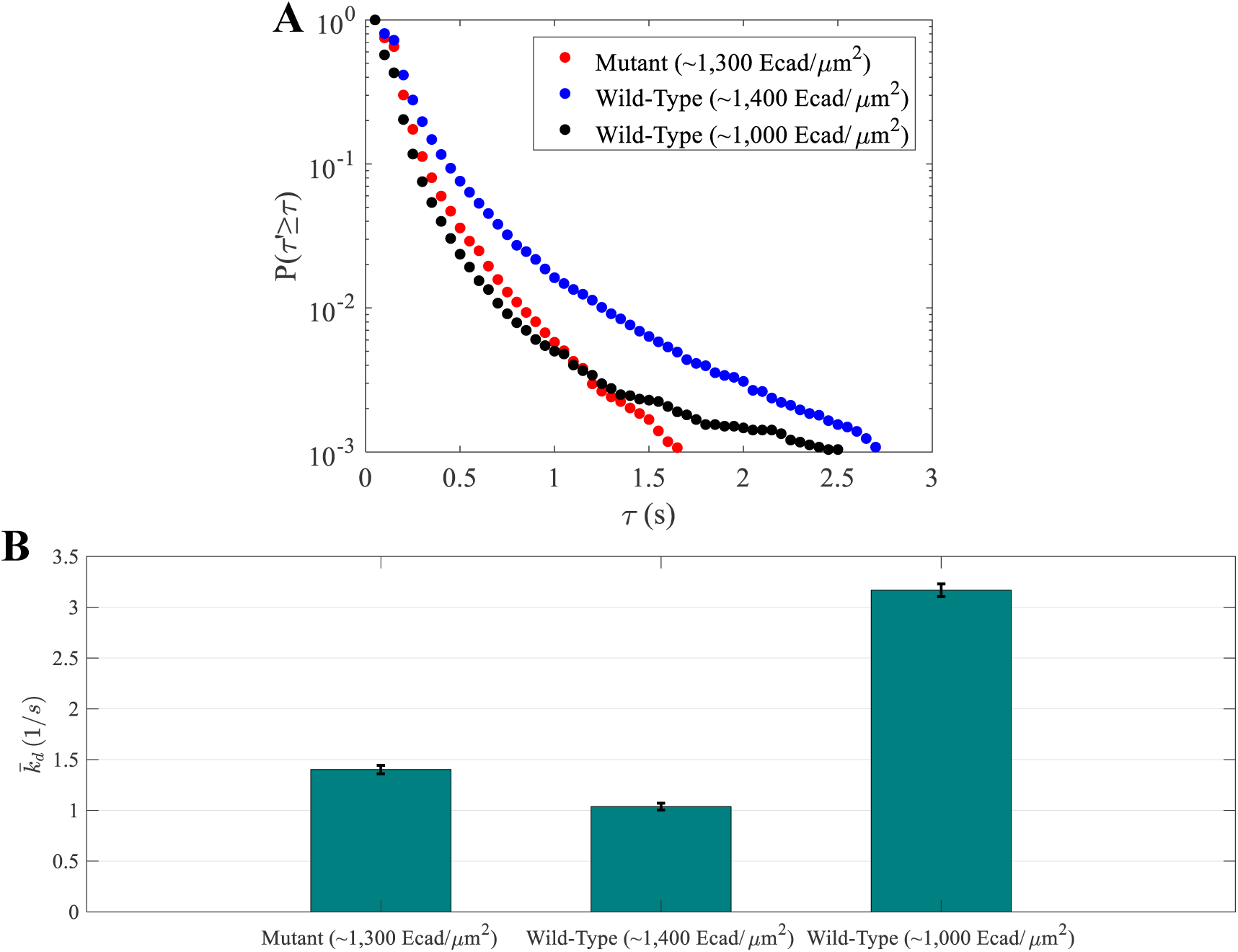
Specific cis-interactions exhibit slow dissociation. (A) High-FRET state complementary cumulative dwell time (i.e. association time) distributions for mutant E-cad and two concentrations of wild-type E-cad. (B) Average dissociation rate constants 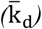 for the mutant and two wild-type conditions resulting from modeling interactions using a Markov model. Error bars were estimated as the square root of the Cramèr-Rao lower bound.

For this system, the distribution of dwell times associated with the high-FRET state corresponds to the time in which a donor E-cad is associated with an acceptor E-cad. However, some of the high-FRET dwell times used to calculate the complementary cumulative dwell time distribution in Fig. 3A are not bounded by transitions to the low-FRET state (e.g., those observed at the beginning or end of trajectories). Consequently, quantitative analysis of the dwell time data is complicated by the fact that the measured dwell times may be coupled to the observation times (i.e. surface residence times) associated with individual molecular trajectories. For example, the probability of observing a long, high-FRET state dwell time is higher within a long trajectory than within a short trajectory. Therefore, the high-FRET state dwell times must in principle be analyzed in conjunction with surface residence times in order to extract quantitative information about protein-protein binding strengths. Qualitatively, since the surface residence time of a high-FRET trajectory is limited by the strength of the lateral protein-protein interactions experienced by that molecule, as opposed to the surface binding affinity or photobleaching (see Supplementary File 1), stronger lateral binding interactions result in both longer associations and longer surface residence times, and the general trends exhibited by the high-FRET dwell time distributions (Fig. 3A) do in fact reflect real differences in the lateral binding affinity. However, in order to rigorously extract quantitative dissociation rates, it was necessary to employ a three-state Markov model that accounted for unbounded dwell times, as described in detail below.

As indicated in Fig. 3A, which shows the association time distributions in the high-FRET state, the probability of a long association time (τ) is significantly higher for the high surface coverage wild-type condition than for the mutant at high coverage and for the wild-type at intermediate surface coverage. Notably, both wild-type distributions (at high and intermediate surface coverages) exhibit slowly-decaying “tails” of long-lasting associations times, suggesting the presence of a binding mode associated with strong protein-protein interactions. This is presumably due to relatively rare, but long-lived, specific interactions through the cis-interface. Consistent with this interpretation, the association time distribution for the cis-mutant E-cad (where specific cis-interactions are absent) lacks this slowly-decaying “tail”, and falls more steeply, consistent with weaker overall protein-protein interactions.

Interestingly, large numbers of short association time intervals are observed for all three conditions, including the mutant and both wild-type concentrations. The slope of the graph in this region (related to the inverse dissociation rate) is much steeper than that of the long time-interval regime of wild-type E-cad. This behavior suggests the presence of an additional weak interaction besides the specific cis-interactions between wild-type E-cadherins. These additional interactions are hypothesized to represent weak nonspecific interactions between the extracellular domains of either the wild-type or mutant E-cad. Interestingly, by observing the initial slope of the distributions, it appears that the effective strength of these nonspecific interactions increases with increasing surface coverage, consistent with previous observations (Langdon, Kastantin, Walder, & Schwartz, 2014). At higher protein surface coverage, binding avidity within large protein clusters and the prevalence of steric effects such as trapping within cluster interiors increase the effective association time.

As described above, a three-state Markov model that has previously been used to model protein conformations based on intramolecular FRET time series data (Kienle, Falatach, Kaar, & Schwartz, 2018) was used to quantitatively model intermolecular FRET time series data associated with E-cad interactions in this system. This model incorporated three states: high-FRET, low-FRET, and off, where the off-state corresponded to the end of a trajectory due to photobleaching, desorption from the surface, or diffusion out of the field of view. To account for heterogeneity in protein interactions, a beta distribution of state transition probabilities between the high-FRET and low-FRET states was incorporated into the model. This heterogeneity reflects the diversity of local environments, including various cluster sizes, shapes, etc. A maximum likelihood estimate of the beta distribution parameters was iteratively generated based on the sequence of states for each trajectory, and the average interaction rates for transitions from the low-FRET state to the high-FRET state and vice versa were determined. Here, the average interaction rate for transition from the high-FRET state to the low-FRET state was equivalent to the average dissociation rate constant 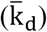 for this system due to the concentration independence of the dissociation reaction rate. For additional details of the model, see the Materials and Methods section and the previous application of this model to protein conformational changes (Kienle et al., 2018). To confirm the accuracy of modeling the observed interactions, complementary cumulative dwell time distributions were generated for comparison with measured distributions, by using the maximum likelihood estimated transition probabilities; they are presented as Fig. S4.

As shown in Fig. 3B, 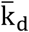 varied significantly between wild-type and mutant E-cad, and also between wild-type E-cad at high and intermediate surface coverage. The values of 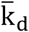 were 1.40 ± 0.04 *s*^−1^, 1.04 ± 0.03 *s*^−1^, and 3.17 ± 0.06 *s*^−1^ for the mutant, at high wild-type surface coverage, and at intermediate wild-type surface coverage, respectively. Thus, wild-type E-cad at high surface coverage exhibited the slowest dissociation (i.e., the most stable clusters), consistent with expectations from the association time distribution. This is plausible, since larger clusters at higher surface concentrations were expected to enable both long-lived multivalent nonspecific interactions as well as a significant number of long-lasting specific cis-interactions. For mutant E-cad at high coverage, the value of 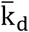 was larger than for wild-type E-cad at high surface coverage, but significantly smaller than for wild-type E-cad at intermediate surface coverage. This was presumably due to the relatively high effective strength of nonspecific interactions, at high surface coverage, due to avidity and trapping effects, as described above. Finally, the largest value of 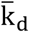 (i.e., the least stable clusters) was observed for wild-type E-cad at intermediate surface coverage, due mainly to the frequent and short-lived nonspecific interactions. This is consistent with previous observations (and the shape of the association lifetime distribution in Fig. 3A) and suggests that specific cis-interactions were infrequent at this intermediate surface coverage.

Overall, an additional interesting result from the modeling of the FRET time-series data was that E-cad interactions were highly heterogeneous under all conditions, presumably due to the wide variety of cluster sizes and shape, the presence of trapping and avidity effects, and the complex combination of specific and nonspecific interactions. The mutant E-cad interactions, which only included nonspecific associations, were particularly heterogeneous; perhaps reflecting the potential for multivalency in these associations. Moreover, the presence of both nonspecific and specific interactions creates many complex scenarios, including the potential for specific cis-interactions to form via an initial nonspecific ‘encounter complex’ that transitions to the specific cis-interaction through orientational changes. To capture this complexity directly, explicit kinetic Monte Carlo simulations were performed, as described below.

### Heterogeneous kMC Simulations Differentiate Specific and Nonspecific Interactions

The single molecule FRET results provided novel insights into the qualitative overall behavior of lateral interactions between E-cad extracellular domains tethered to a supported bilayer. They also enabled quantitative characterization of the dissociation kinetics due to specific and nonspecific interactions. Nevertheless, gaps remained in the understanding of the physical basis of the observations. In particular, as discussed above, it was difficult to unambiguously distinguish association events. Additionally, single molecule FRET permitted the assignment of only two states: low-FRET and high-FRET (associated). Therefore, for a system in which intrinsically different (and highly heterogeneous) interactions are expected, these experimental observations could not distinguish between the different types of interactions underlying clustering. Nor could we quantitatively extract the independent contributions and kinetics of each interaction. To address these experimental limitations, kMC simulations were performed. Importantly, these simulations incorporated both the nonspecific and specific interactions revealed by the FRET data.

To model specific interactions, each wild-type E-cad molecule had one cis-donor site and one cis-acceptor cite located on opposing sides of the molecule (see Fig. 4A), in order to incorporate the specific orientational constraint associated with specific cadherin cis-interactions (Oliver J. Harrison et al., 2011). This allowed each E-cad molecule to participate in a maximum of two specific cis-interactions, and mandated the formation of flexible linear oligomers. The inclusion of nonspecific interactions was then accomplished by allowing additional interactions in all directions, within a specified distance constraint. By allowing molecules to form both nonspecific and specific interactions, association and dissociation rate constants could be tuned independently for both interactions.

**Fig. 4.**
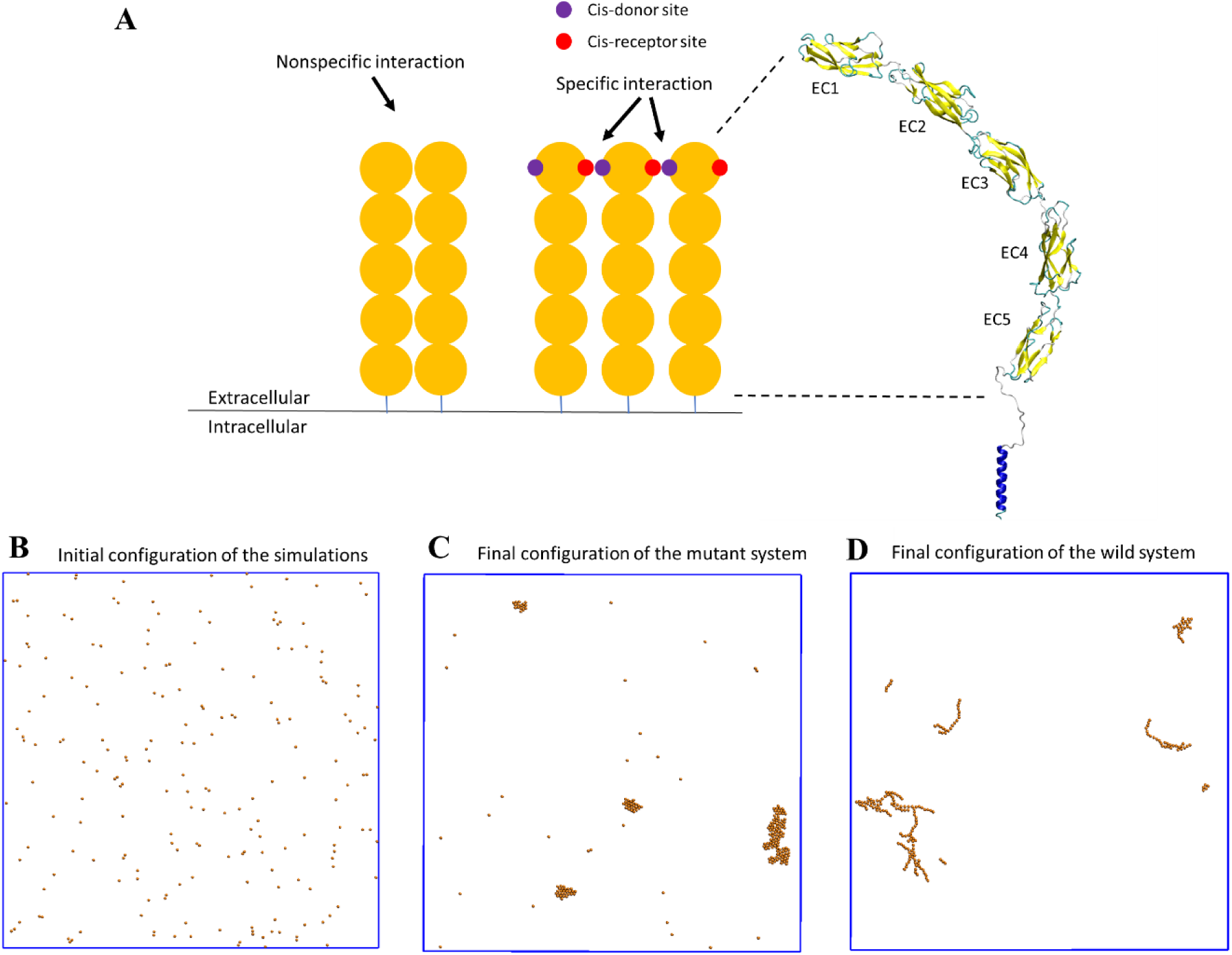
A coarse-grained model was constructed to simulate the spatial-temporal process of E-cad clustering. (A) E-cad extracellular domains (orange), nonspecific and specific cis-interactions. Cis-donor sites are labeled in purple, and cis-receptor sites are labeled in red. A structural model of the E-cad is shown on the right side. Ectodomain structure with EC domains 1-5 numbered from the N-terminus. (B) Top view of initial configuration in the simulations. The number of E-cad molecules is equal to 200. (C) Top view of final configuration in the mutant system. (D) Top view of final configuration in the wild-type system.

We computationally simulated the clustering of E-cad on supported lipid-bilayers, using a domain-based, coarse-grained model (Fig. 4A). After random initial placement, all molecules and clusters stochastically diffused off-lattice, using periodic boundary conditions. The average cluster size was monitored throughout the simulation period. Simulations were run until the average cluster size did not change significantly. This implied that equilibrium was reached, analogous to the experiments. A total of 50 simulations were run at three different surface coverages (312.5 E-cad/µm^2^, 625 E-cad/µm^2^, and 1,250 E-cad/µm^2^) for both wild-type and mutant E-cad. Simulations with wild-type E-cad included both nonspecific and specific interactions, but simulations of cis-mutants allowed the proteins to associate only by nonspecific interactions. Simulations also used different combinations of binding rates within a biologically relevant range. For additional details on kMC simulations, see the Materials and Methods section.

To model specific interactions, each wild-type E-cad molecule had one cis-donor site and one cis-acceptor cite located on opposing sides of the molecule (see Fig. 4A), in order to incorporate the specific orientational constraint associated with specific cadherin cis-interactions (Oliver J. Harrison et al., 2011). This allowed each E-cad molecule to participate in a maximum of two specific cis-interactions, and mandated the formation of flexible linear oligomers. The inclusion of nonspecific interactions was then accomplished by allowing additional interactions in all directions, within a specified distance constraint. By allowing molecules to form both nonspecific and specific interactions, association and dissociation rate constants could be tuned independently for both interactions.

We computationally simulated the clustering of E-cad on supported lipid-bilayers, using a domain-based, coarse-grained model (Fig. 4A). After random initial placement, all molecules and clusters stochastically diffused off-lattice, using periodic boundary conditions. The average cluster size was monitored throughout the simulation period. Simulations were run until the average cluster size did not change significantly. This implied that equilibrium was reached, analogous to the experiments. A total of 50 simulations were run at three different surface coverages (312.5 E-cad/µm^2^, 625 E-cad/µm^2^, and 1,250 E-cad/µm^2^) for both wild-type and mutant E-cad. Simulations with wild-type E-cad included both nonspecific and specific interactions, but simulations of cis-mutants allowed the proteins to associate only by nonspecific interactions. Simulations also used different combinations of binding rates within a biologically relevant range. For additional details on kMC simulations, see the Materials and Methods section.

For each set of simulation parameters, multiple independent trajectories were generated to assure that the computational data were statistically meaningful. Detailed strategies of the sensitivity analysis are summarized in the Materials and Methods section. At the end of the simulations, the cluster size distributions were calculated by averaging from all the trajectories in the systems. In order to directly compare the cluster size distributions from simulations with the experimental distributions, similar surface coverages were considered between the simulation and experimental systems.

To allow direct comparison of kMC simulations to experimental results, E-cad cluster size probability distributions were calculated using raw trajectory friction factor data adapted from Thompson, et al. (Thompson et al., 2019), as described in the Materials and Methods section. Resulting experimental cluster size probability distributions are shown below as Fig. 5A-B, for both wild-type and mutant E-cad at high, intermediate, and low surface coverages corresponding to ∼39,000 E-cad/µm^2^, ∼1,000 E-cad/µm^2^, and ∼0.6 E-cad/µm^2^, respectively. For mutant E-cad, the change in the cluster size distribution with increasing surface coverage is subtle, and mainly visible in the small cluster regime, where the peak present at low surface coverage at ∼20 E-cad shifts to a modestly larger cluster size of ∼40 E-cad. This change is presumably due to weak nonspecific interactions between the mutants that support cluster formation at elevated surface coverage. The cluster size distributions of wild-type E-cad exhibit a more dramatic change with increasing surface coverage, particularly in the tails of the distributions. For example, at high and intermediate surface coverage the probability of observing a large cluster (∼40 to ∼160 E-cad) is significantly increased. This change with increasing surface coverage for wild-type E-cad is likely due to a combination of nonspecific and specific interactions causing large cluster formation, relative to the cluster formation observed for the mutant.

**Fig. 5.**
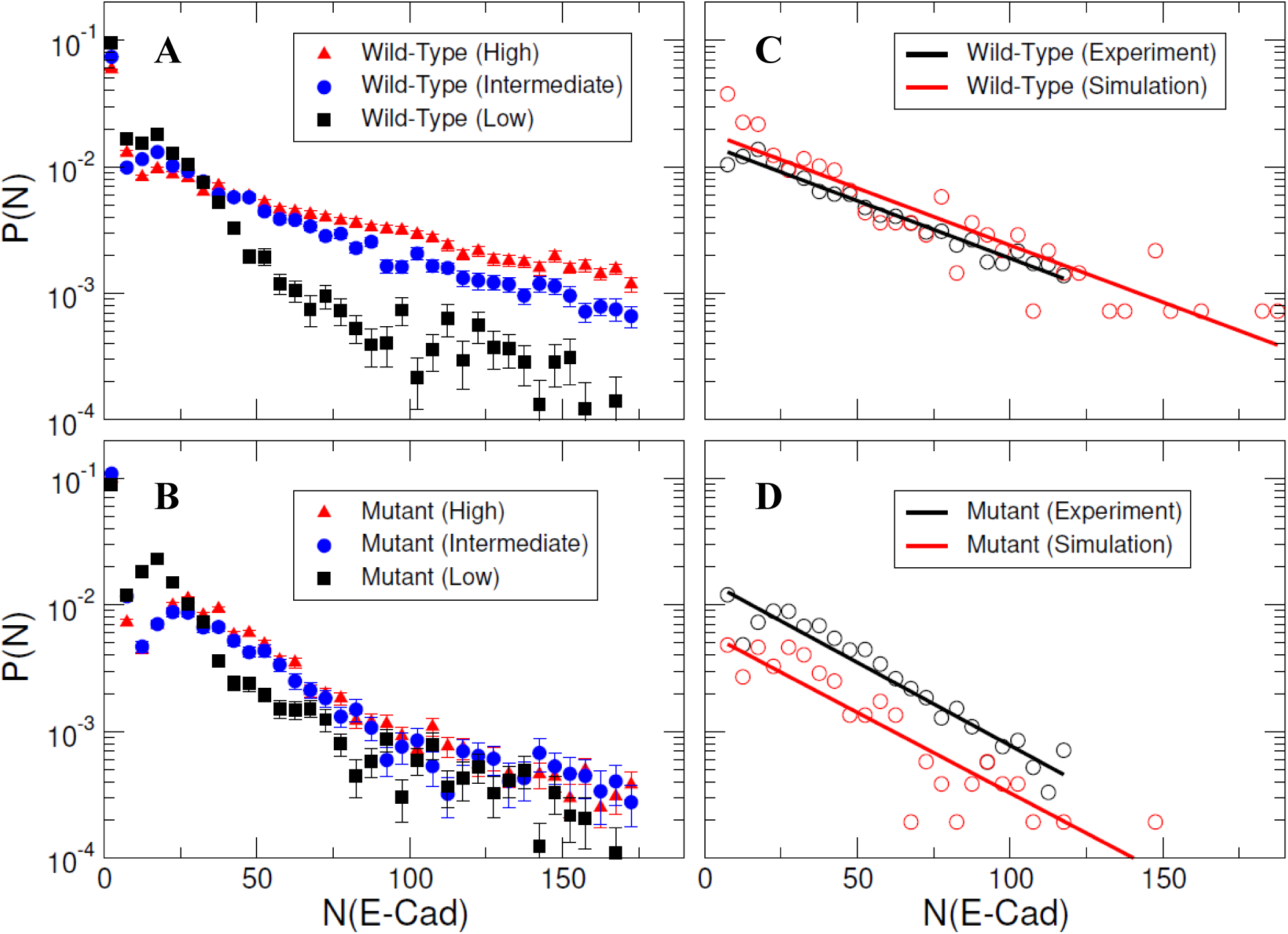
Specific and nonspecific interactions can cause E-cad clustering. (A-B) Representative experimental cluster size probability distribution functions for wild-type and mutant E-cad at low, intermediate, and high surface coverages. Error bars correspond to the standard deviation of cluster size probability distribution functions calculated using 100 samples using a bootstrap method with replacement. (C-D) The comparison of experimental and simulated cluster size distributions for mutant and wild-type E-cad. The solid lines indicate the single exponential fitting.

For kMC simulations, we first turn off specific cis-interactions, so that E-cad can form clusters only through the nonspecific lateral interaction. This simulation is used to mimic the system in which the mutant is employed to eliminate specific cis-interactions. The final configuration from a representative simulated trajectory is shown in Fig. 4C. In addition to E-cad monomers, homogeneously distributed compact clusters formed through nonspecific cis-interactions between mutant E-cad proteins. Fig. S8 further shows the cluster size distributions under different on/off rate combinations of the nonspecific interactions. Cluster size distributions can be fitted by a single exponential function 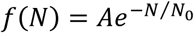 where N_0_ corresponds to the characteristic cluster size. Fig. S8 indicates that the characteristic cluster size is closely related to the values of the on and off rates. The simulated on and off rates were therefore optimized so that the cluster size distribution from simulations (red) agreed with the experimental distribution (black) for the cis-mutant (Fig. 5D). The value of the characteristic cluster size in the experiment was ∼29 E-cad, which is equal to the computational characteristic cluster size of ∼29 E-cad, within experimental uncertainty. The on and off rates of the nonspecific interaction used to generate the distribution in the simulation are 2×10^5^ s^−1^ and 10^3^ s^−1^, respectively (Table S8). These on/off *rates* correspond to the effective *rate constants* of *k*_*on*_≅1.1×10^6^ M^−1^ s^−1^ and *k*_*off*_≅1×10^3^ s^−1^, based on the calculation developed in our previous studies (Wang, Xie, Chen, & Wu, 2018). These rates correspond to an effective binding affinity in the mM range, for nonspecific cis-interactions.

Subsequently, we carried out simulations in which the specific cis-interaction was turned on. Different combinations of on/off rates for the specific interaction were systematically tested, while the rates of the nonspecific interactions were fixed at the values determined for the cis-mutant. The final configuration from one of these simulations is shown in Fig. 4D. Relative to the homogeneous and compact clusters observed in the simulations associated with E-cad mutant, the clusters formed when both nonspecific and specific cis-interactions were switched on exhibited extended (linear) configurations. These one-dimensional linear clusters are derived from the polarized cis-binding interface, which is inferred from the x-ray crystal structure of wild-type E-cad (Oliver J. Harrison et al., 2011). Cluster size distributions associated with different combinations of on and off rates for specific interactions are shown in Fig. S9. Again, we identified an appropriate combination of specific cis on/off rates that resulted in a similar characteristic cluster size as was observed experimentally for wild-type E-cad, as shown in Fig. 5C. The value of the characteristic cluster size for the experiment is ∼33 E-cad, which is very similar to the computational value of ∼34 E-cad from simulations. The on and off rates of the specific interaction that were used to generate the distribution in the simulation are 10^8^ s^−1^ and 10^2^ s^−1^, respectively (Table S8). These on/off *rates* for the specific cis-interaction correspond to the effective *rate constants* of *k*_*on*_≅2.7×10^6^ M^−1^s^−1^ and *k*_*off*_≅1×10^2^ s^−1^, and to a binding affinity of approximately 10 µM. Comparisons of the specific and nonspecific interactions suggest that the specific cis-binding rate is slightly faster than that of the nonspecific interaction, and the specific cis-interaction is stronger by approximately an order of magnitude.

Finally, in addition to comparisons of cluster size distributions, association time distributions extracted from the simulations were also calculated and qualitatively compared to the experimental association time distributions discussed in the previous section. This ensured that the simulations captured the experimental behavior. Fig. 6 shows the comparison of experimental and simulated association time distributions for mutant and wild-type E-cad. In both simulations and experimental measurements, the association time of E-cad increases when in the presence of specific cis-interactions (wild-type vs. cis-mutant), demonstrating qualitative consistency. We note that the dwell-time distributions from simulations are not necessarily expected to agree quantitatively with experimental measurements, due in part to the difference between the experimental acquisition time (50ms) and simulation time step (0.01ns). Notably, the long-time asymptotic behavior of experimental and simulated dwell times have similar behavior (i.e. the slopes of the distribution tails in Fig. 6), indicating that the simulations accurately capture the salient experimental behavior. Furthermore, experimental phenomena such as desorption, photobleaching, and supported lipid bilayer defects and heterogeneity are not accounted for in the simulations and may limit quantitative comparisons of association times. Overall, these simulation results are qualitatively consistent with longer-lived wild-type E-cad interactions. This is due to specific cis-interactions, as well as to the potential interplay between nonspecific and specific interactions.

**Fig. 6.**
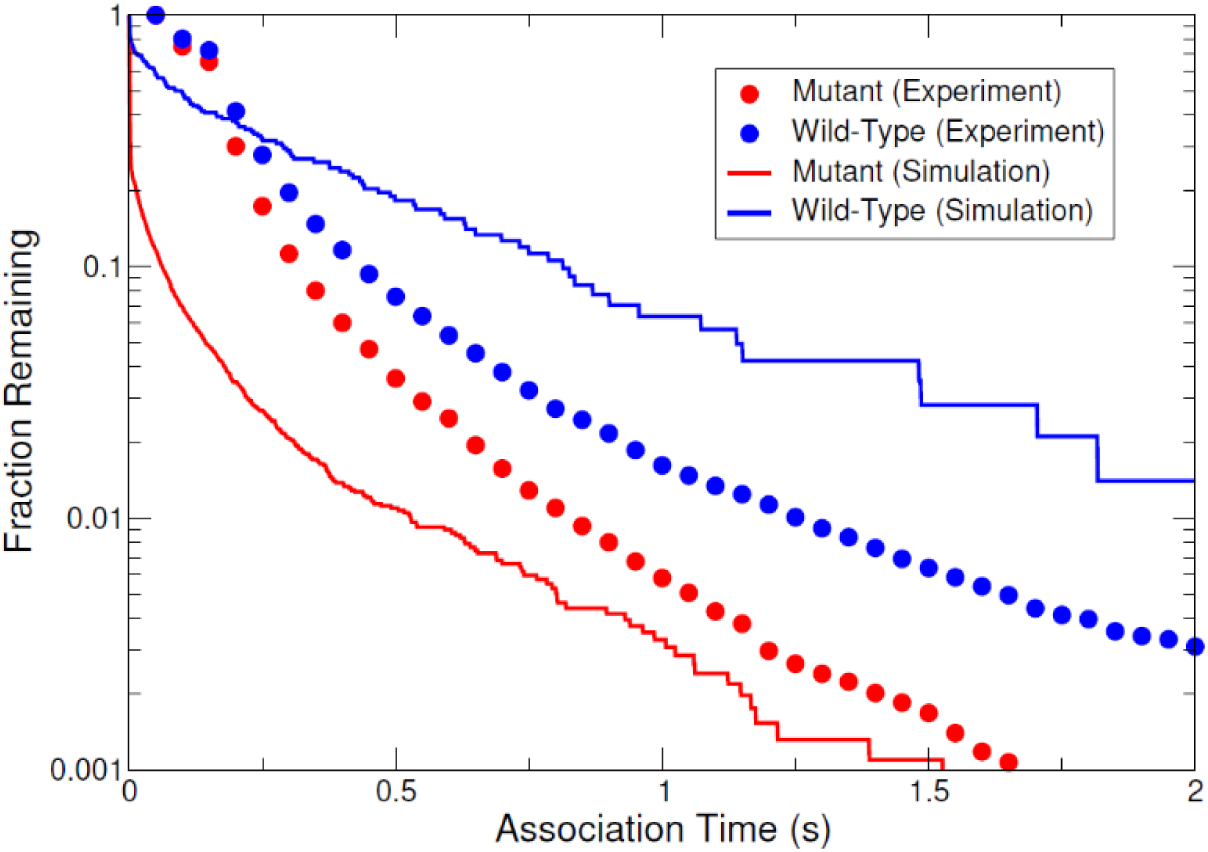
The comparison of experimental and simulated complementary cumulative association time distributions for mutant and wild-type E-cad.

## Discussion

An important advance of this research involves the development of a combined experimental and theoretical framework that enables the quantification of lateral binding interactions between proteins confined to fluid, 2D membrane bilayers. The single molecule FRET measurements revealed that both specific and nonspecific cis-interactions contribute to wild-type E-cadherin clustering at a physiologically relevant surface coverage. Complementary kMC simulations provided important insights into the molecular events underlying the FRET distributions, and further extracted rate constants for both specific and nonspecific lateral interactions between the cadherin extracellular domains. Moreover, these results successfully demonstrated directly that E-cadherin extracellular domains associate through cis-interactions. Prior experimental data supported the role of specific cis-interactions in the assembly of cadherin clusters, both at intercellular adhesions and on supported lipid bilayers at high surface densities. However, until recently, direct characterization of E-cad cis-interactions was not possible by traditional methods, due to the weak binding affinity.

Notably, we find that both specific and nonspecific interactions control E-cad clustering on membranes at high surface coverage, and that nonspecific interactions contribute to both mutant E-cad and wild-type E-cad lateral interactions at surface concentrations below the surface coverage threshold for cis-clustering. Although these nonspecific interactions are weaker than specific cis-interactions, they are more frequent, and hence dominate at low concentrations. The conditions employed in these measurements isolated the effects of specific and nonspecific interactions, and enabled quantitative comparisons with kMC simulations. For both the mutant and wild-type E-cadherin at intermediate surface coverage, where the intermolecular interactions are primarily due to nonspecific interactions, the high-FRET state corresponds to slower diffusion than the low-FRET state. The latter behavior is a result of small, short lived, cluster formation, and was only observable due to the ability to isolate high-FRET objects. However, if one were only able to compare the overall average diffusion of all objects, the slight decrease in the diffusion coefficient of mutant E-cad at high concentration would not be observable, as previously reported (Thompson et al., 2019).

It was necessary to include both nonspecific and specific interactions in the kMC simulations, in order to accurately reproduce the experimental cluster size distributions. This agreement confirmed the interpretation of the single-molecule FRET data. The rate constants associated with each of these distinct lateral interactions further show that, despite the 10-fold slower dissociation rate of specific cis-bonds, the nonspecific interactions must be taken into account.

The influence of nonspecific interactions on mutant E-cad has not previously been reported. Indeed, it was necessary to combine highly sensitive single-molecule FRET with computational simulations, and to explicitly compare wild-type and cis-mutant E-cad, in order to characterize these weak interactions. Moreover, as these results demonstrate, nonspecific interactions are dynamic and short lived, and would not likely be detected by alternative methods, such as ensemble averaged FRET or photon counting (Biswas et al., 2015; Zhang et al., 2009). Although nonspecific steric (repulsive) interactions have been invoked to account for membrane protein organization (Albersdörfer, Feder, & Sackmann, 1997; Paszek et al., 2014; Qi et al., 2001; Schmid et al., 2016), the potential significance of nonspecific attractive interactions was not fully appreciated prior to this study.

E-cadherin represents a special, and particularly demanding test case for characterizing lateral protein interactions tethered to lipid bilayers, because the cis-bonds have very low affinity and are not detectable in solution. This combination of single molecule FRET and kMC simulations can be extended to other proteins such as nectins that likely interact through higher affinity cis-bonds (Rikitake et al., 2012). Although there are approaches for quantifying the 2D trans- (adhesive) affinities and binding rates of membrane receptors, until now, few measurements were able to quantify lateral binding affinities (Chen, Novicky, Merzlyakov, Hristov, & Hristova, 2010; Chesla et al., 1998; Chien et al., 2008; Sarabipour, Del Piccolo, & Hristova, 2015; J. Wu et al., 2008; D. M. Zhu, Dustin, Cairo, & Golan, 2007), and there are no prior reports of measured binding rates. Interestingly, theoretical models of cadherin binding predict cooperativity between trans-binding between opposing cadherins and cis-interactions (Y. Wu et al., 2010). The approach described in this study lays the groundwork for directly testing that hypothesis, by comparing cis-binding rates, for example, between cadherins on free membranes versus within adhesion zones.

These findings provided new insights regarding the physical interactions underlying E-cadherin clustering. They also raise the possibility that nonspecific interactions could similarly influence the oligomerization of other membrane proteins. Conversely, the methods described in this study also open the possibility of quantifying the impact of other factors such as crowding, confinement, or even membrane topography on protein interactions.

## Materials and Methods

### FRET Sample Preparation

CEP 4.2 plasmids encoding the hexahistidine-tagged wild-type E-cad and L175D mutant were obtained from Dr. Lawrence Shapiro (Columbia University, NY). The Human Embryonic Kidney 293T (HEK293T) cell line was from the American Type Culture Collection (Manassas, VA). Cells were cultured in Dulbecco’s Minimum Eagle Medium (DMEM) containing 10% fetal bovine serum (FBS) (Life Technologies, Carlsbad, CA) under 5% CO_2_ atmosphere at 37 °C. Cell lines that stably expressed the soluble proteins were generated, by transfecting HEK293T cells with the mutant construct, using Lipofectamine 2000 (Invitrogen, Grand Island, NY) according to the manufacturer’s instructions.

HEK293T cell lines that stably expressed hexahistidine-tagged, soluble E-cadherin ectodomains were selected with 200 µg/mL Hygromycin B (Invitrogen). Western blots of the culture medium confirmed protein expression by individual colonies. The colonies that expressed the highest levels of soluble protein were pooled for further protein production. Secreted, hexahistidine-tagged cadherin was then purified from filtered culture medium, by affinity chromatography with an Affigel NTA affinity column, followed by ion-exchange chromatography (Aktapure). Protein purity was assessed by SDS polyacrylamide gel electrophoresis, and adhesive function was confirmed with bead aggregation assays (Sivasankar, Brieher, Lavrik, Gumbiner, & Leckband, 1999).

Purified E-cad extracellular domains with C-terminal 6xHis tags were randomly labeled using an Alexa Fluor 555 (AF555) NHS-ester antibody labeling kit, and both wild-type and L175D mutant were labeled using an Alexa 647 (AF647) NHS-ester antibody labeling kit (succinimidyl ester; Invitrogen, Carlsbad, CA). Protein was reacted with the dye for 1 h in buffer (25 mM HEPES, 100 mM NaCl, 10 mM KCl, 2 mM CaCl_2_, 0.05 mM NiSO_4_, pH 8) at room temperature. Unreacted dye was removed via spin column. Based on absorbance measurements, using extinction coefficients of 150,000 cm^−1^ M^−1^ for the AF555, 239,000 cm^−1^ M^−1^ for the AF647, and 59,860 cm^−1^ M^−1^ for the protein, the labeling stoichiometry was ∼1.3 for AF555 labeling of wild-type E-cad and ∼2.3 and ∼1.3 for AF647 labeling of wild-type and mutant E-cad, respectively.

1,2-Dioleoyl-sn-glycero-3-phosphocholine (DOPC) was purchased from Millipore Sigma (Burlington, MA). 1,2-dioleoyl-sn-glycero-3-[(N-(5-amino-1-carboxypentyl)iminodiacetic acid)succinyl] (nickel salt) (DGS-NTA(Ni)) was purchased from Avanti Polar Lipids (Alabaster, Alabama). DOPC and DGS-NTA(Ni) were dissolved in chloroform in the molar ratio of 19:1 in a glass culture tube. Following solvent evaporation under a stream of nitrogen, a thin film of lipids was formed on the side of the tube. This lipid film was then hydrated with buffer so the total lipid concentration was 3 mM. This suspension was mixed via vortex and sonicated for 0.5 h. The vesicles were then extruded through a 50 nm filter membrane (Whatman, Maidstone, UK) 21 times to form unilamellar vesicles with a homogeneous size distribution.

Glass coverslips (Fisher Scientific, Hampton, NH) and fused silica wafers (Mark Optics, Santa Ana, CA) were cleaned with piranha solution for 2 h and treated by UV-ozone for 0.25 h. Following surface treatment, the wafers were placed in a custom built flow cell that had been cleaned using Micro-90 detergent solution (International Product Corp., Burlington, NJ). To form supported lipid bilayers, a dispersion of unilamellar vesicles (3 mM total lipid concentration) was carefully injected into the flow cell in order to avoid air bubble formation. Following a 1 h incubation period, vesicles spontaneously formed a supported lipid bilayer via vesicle fusion (Cremer & Boxer, 1999; Richter, Berat, & Brisson, 2006). Following formation, the bilayer was rinsed with buffer to remove excess vesicles and incubated with 100 mM NiSO_4_ for 0.5 h to ensure complete chelation of DGS-NTA(Ni) lipids. The supported lipid bilayer was then exchanged into buffer before injecting 300 µL of a protein buffer solution containing AF555 labeled wild-type E-cad and either AF647 labeled wild-type E-cad and unlabeled wild-type E-cad or AF647 labeled mutant E-cad and unlabeled mutant E-cad, permitting the binding of hexahistidine-tagged E-cad to the DGS-NTA lipids. In this configuration, the AF555 labeled E-cad served as the FRET donor and the AF647 labeled E-cad served as the FRET acceptor. Two different total wild-type E-cad solution concentrations of 3×10^−7^ M and 5×10^−7^ M and one total mutant E-cad solution concentration of 5×10^−7^ M were studied. Table S3 summarizes the donor and acceptor solution concentrations for the three conditions. The donor concentration was adjusted to allow for single molecule resolution, and the acceptor concentration was optimized to allow for a large number of FRET events, but an insignificant amount of direct excitation of the acceptor. Using the optimized donor and acceptor concentrations, donor bleed through into the acceptor channel and direct excitation of the acceptor were both determined to be insignificant by imaging control samples containing either donor and unlabeled E-cad or acceptor and unlabeled E-cad and checking for significant emission in the acceptor channel. The addition of unlabeled E-cad was necessary in order to reach a surface coverage high enough, such that significant cluster formation had occurred (Thompson et al., 2019). This resulted in a large number of high-FRET events, indicated by an acceptor intensity greater than that of the donor. This high surface coverage could not be achieved by only adding donor and acceptor E-cad, due to overwhelming emission in the acceptor channel due to direct excitation of the acceptor.

### Single-Molecule TIRFM FRET Imaging

Imaging of the samples was accomplished using a custom-built prism-based TIRF microscope (Nikon TE-2000 base, 60x water-immersion objective, Nikon, Melville, NY). Custom-built flow cells were mounted on the microscope stage and a 532 nm 50 mW diode-pumped solid state laser (Samba, Cobolt, Solna, Sweden) was used as an excitation source, incident through a hemispherical prism in contact with the wafer on the top of the flow cell. This resulted in an exponentially decaying TIRF field propagating into solution, selectively exciting donor fluorophores at the lipid bilayer-water interface. Fluorescent emissions from the donor and acceptor were separated using an Optosplit III beam splitter (Cairn Research, Faversham, UK) containing a dichroic mirror with a separation wavelength of 610 nm (Chroma, Bellows Falls, VT). Fluorescence from the donor and acceptor were further filtered using a 585/29 bandpass filter and 685/40 bandpass filter (Semrock, Rochester, NY), respectively. The donor and acceptor channels were then projected onto different regions of an Andor iXon3 888 EMCCD camera (Oxford Instruments, Abingdon, UK) maintained at −95 °C. An acquisition time of 50 ms was used to capture 12 or 13 image sequences (i.e. movies) of each sample, where each movie was 5 min long. Additionally, to allow for accurate donor and acceptor colocalization, the donor and acceptor channels were aligned using images of a glass slide that had been scratched with sand paper, resulting in an irregular alignment image. The details of this image alignment process are described previously (Faulón Marruecos, Kienle, Kaar, & Schwartz, 2018).

### Image Analysis

All two-channel movie analysis was performed using custom Matlab-based software, where the methods for determining object positions and intensities and linking trajectories have been described elsewhere (Faulón Marruecos et al., 2018). To briefly summarize, objects that were detected in consecutive frames that were within a user-defined tracking radius (3 pixels or 1.29 µm, for this analysis) were linked into trajectories that could be further analyzed. Object identification was determined using an automated thresholding function that has been described previously (Kienle & Schwartz, 2019). This automatic thresholding software allowed for a user-defined number of noise-objects per frame to be identified, as well as the use of a user-defined object radius (0.05 and 1 pixel for this work, respectively). Objects that were identified within 2 pixels in separate channels were identified as a donor-acceptor pair undergoing FRET. The position of the FRET pair was determined using the object with the greatest signal-to-noise ratio. The FRET state of each object at every frame was assigned using a method and algorithm described elsewhere (Chaparro Sosa et al., 2018). To summarize, two-dimensional heat maps showing the donor intensity (*I*_*D*_) versus acceptor intensity (*I*_*A*_) were constructed. It was apparent that two populations were present at high and low FRET efficiency. A linear threshold dividing these two populations was calculated by determining the slope and intercept that minimized the integrated heat map values along the dividing line.

Additionally, due to a combination of bright contaminants and inherent defects in supported lipid bilayers, a permanently immobile (or highly confined) population was observed in the donor channel, which was readily distinguished from the mobile population of interest (Knight, Lerner, Marcano-Velazquez, Pastor, & Falke, 2010). To avoid inclusion of these anomalous trajectories, only trajectories with a median donor intensity less than the 60^th^ percentile were analyzed further, effectively excluding the tail of the median donor intensity distribution. For short-time diffusion coefficient determination, only trajectories with a total surface residence time of at least 0.71 s were included, to allow for significant statistical analysis. This surface residence time minimum of 0.71 s was not required for the dwell time distribution calculations and rate estimations. Therefore, all trajectories longer than 0.1 s (2 frames) were included.

### Surface Coverage Estimation

The surface coverage in terms of # of E-cad/µm^2^ was estimated according to:

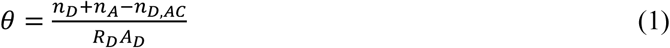

where *θ* is the surface coverage in terms of # of E-cad/µm^2^, *n*_*D*_ is the number of fluorescent molecules in the donor channel, *n*_*A*_ is the number of fluorescent molecules in the donor channel, *n*_*AC,D*_ is the apparent number of fluorescent molecules in the donor channel for an acceptor control sample that did not contain any donor labeled E-cad, *A*_*D*_ is the area of the donor channel, and *R*_*D*_ is the ratio of donor-labeled protein to total protein. Subtracting the apparent number of fluorescent molecules in the donor channel for a sample without any donor labeled E-cad allowed for the exclusion of contamination in the donor channel, as well as any fluorescence in the donor channel from direct excitation of the acceptor. This estimate is assuming a one-to-one transfer of energy from donor to acceptor, complete transfer of energy from donor to acceptor, and minimal apparent objects in the acceptor channel that were not actually FRET acceptors. These assumptions were appropriate for these experiments, primarily because the number of objects in the donor channel was much greater than the number of objects in the acceptor channel and because intermediate FRET-states were not significant. Even so, the resulting surface coverage values should be treated as estimates. The fractional surface coverage was averaged over only the first ten frames of each movie to minimize the underestimation of surface coverage due to photobleaching. To further improve estimates, only objects that were tracked for three frames or more were included in surface coverage calculations. This greatly reduced the inclusion of false noise objects that were observed only for one or two frames. These surface coverage values were converted to a fractional areal surface coverage by multiplying by the cross-sectional area of an E-cad extracellular domain, ∼9 nm^2^, assuming the proteins were in an extended conformation due to the presence of calcium (Lambert et al., 2005; Nagar, Overduin, Ikura, & Rini, 1996). Surface coverage estimates are shown in Table S4, both in terms of # of E-cad/µm^2^ and fractional surface coverage by area, for the three protein solution conditions.

### Average Short-Time Diffusion Coefficient Determination

All molecular displacements between consecutive frames were separated based on FRET state, and complementary cumulative squared displacement distributions were calculated using histograms of all squared displacements in each of the two states (high and low FRET efficiency), where the squared displacement was defined as the square of the Euclidean distance traveled from frame to frame. Additional distributions were constructed using all molecular displacements from both FRET-states. These distributions were then fitted to a Gaussian mixture model:

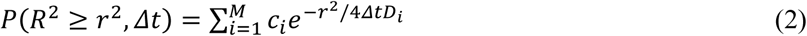

where *r* is the Euclidean displacement between frames, *Δt* is the time between frames (0.05 s), *c*_*i*_ is the fraction of displacements fitted by the *i*th Gaussian term, *D*_*i*_ is the diffusion coefficient for the *i*th term, and *M* is the number of terms included in the model. These data were satisfactorily modeled by *M* = 3. Using the Gaussian mixture model parameters determined from nonlinear fitting, an average short-time diffusion coefficient 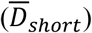 was calculated for both FRET-states and overall:

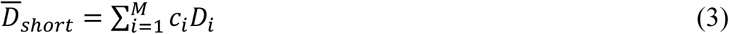

where 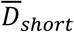 represented the average diffusion coefficient on the shortest experimentally accessible time-scale.

### Surface Residence Time Distributions

Complementary cumulative residence time distributions were constructed for both the high and low-FRET states by separating all trajectories into high-FRET and low-FRET trajectories, where a high-FRET trajectory was defined as any trajectory where the molecule was in the high-FRET state for at least one frame. After trajectory classification, the fraction of molecules that remained on the bilayer a given time after their initial observation (*t*_*s*_) was calculated for both high-FRET and low-FRET trajectories.

### FRET-State Dwell Time Distributions and Transition Rate Determination

Complementary cumulative dwell time distributions were calculated for the two FRET states, corresponding to high and low FRET efficiency, where the apparent dwell time (*τ*) was defined as the number of consecutive frames a trajectory spent in a given state multiplied by the acquisition time, where the FRET state was determined as described above. For these distributions, all dwell times were used, not only dwell times bounded by transitions. This allowed for a large increase in useable data, as many trajectories did not experience multiple FRET state transitions.

Furthermore, E-cad interactions were modeled using a 3-state Markov model that has been previously used to model protein conformation changes (Kienle et al., 2018). To summarize, this model allowed for three states: high-FRET, low-FRET, or off. Therefore, the transition probability matrix had the form:

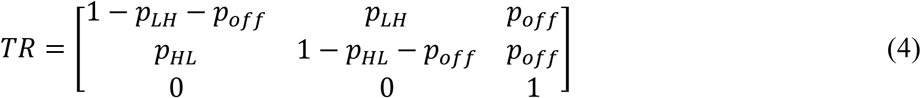

Where *p*_*LH*_, *p*_*HL*_, and *p*_*off*_ are the probabilities for a transition from the low-FRET state to the high-FRET state, from the high-FRET state to the low-FRET state, and for a trajectory to terminate via photobleaching or desorption, respectively. The value of *p*_*off*_ was determined independently by fitting the surface residence times to an exponential distribution. In order to determine the transition probabilities, a maximum likelihood estimate was used based on all trajectory state sequences. To describe the heterogeneity in these transition probabilities, a likelihood function was defined to allow for beta-distributed transition probabilities. The resulting likelihood function was:

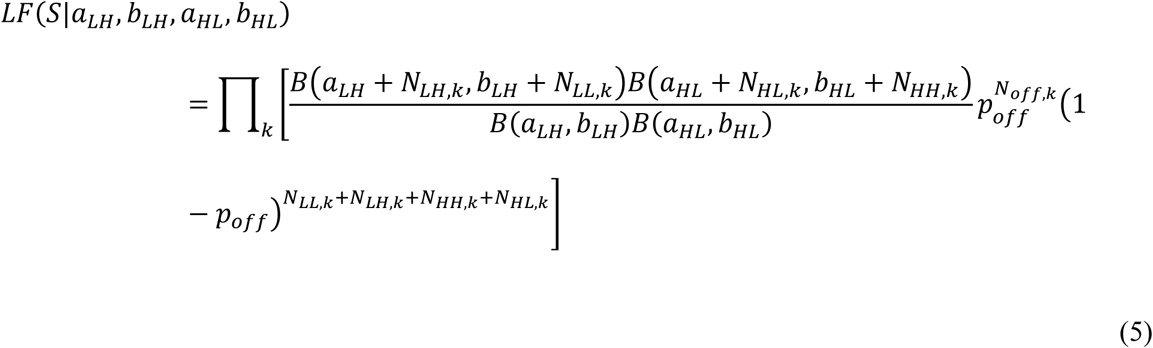

Where *S* is the sequence of observed FRET states for the *k*th trajectory, B is the beta function, and *N*_*HL,k*_, *N*_*LH,k*_, *N*_*LL,k*_, *N*_*HH,k*_, and *N*_*off,k*_ are the number of times within the *k*th trajectory the molecule transitions from the high-FRET state to the low-FRET state, transitions from low-FRET state to the high-FRET state, remains in the low-FRET state, remains in the high-FRET state, and ends, respectively. The model is parameterized by *a*_*LH*_, *b*_*LH*_, *a*_*HL*_, and *b*_*HL*_, which are the parameters defining the beta distribution of *p*_*LH*_ and *p*_*HL*_, respectively. The log of this likelihood function was maximized by iteratively changing the parameters defining the beta distributions describing the transition probabilities between the high and low-FRET states. The average transition rates were then estimated by:

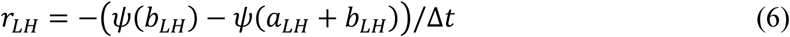

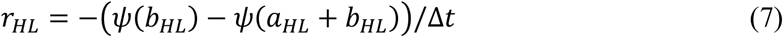

where Δ*t* is the experimental acquisition time, *ψ* is the digamma function, and *r*_*LH*_ and *r*_*HL*_ are the average transition rates from the low-FRET state to the high-FRET state and from the high-FRET state to the low-FRET state, respectively. Additionally, for transition from the high-FRET state to the low-FRET state, the average transition rate is equivalent to the average dissociation rate constant 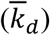, since dissociation is a unimolecular reaction. This is not the case for transition from the low-FRET state to the high-FRET state. Resulting beta distributions of state transition probabilities are shown as Fig. S5, and the corresponding probability density functions for state transition rates are shown as Fig. S6. The values of the average transition rates are presented as Table S5. After determining the most likely beta distribution parameters for the transition probabilities, trajectories were simulated using these transition probability distributions and complementary cumulative dwell time distributions were constructed after truncating the simulated trajectories by sampling from the experimental trajectory surface residence time distributions. These theoretical dwell time distributions were compared to the experimental distributions to check for model consistency (Fig. S4).

### Single-Molecule TIRFM Cluster Size Distributions

In order to calculate E-cad cluster size distributions, raw trajectory friction factor data were adapted from Thompson, et al. and subjected to further analysis (Thompson et al., 2019). Briefly describing the methods used to generate these raw friction factor data: TIRFM was used to observe single AF555 labeled E-cad molecules diffusing on DOPC supported lipid bilayers containing 5% DGS-NTA(Ni) as a function of increasing E-cad surface coverage. Single molecule trajectories were extracted and an effective diffusion coefficient (*D*_*T*_) was calculated for each trajectory according to:

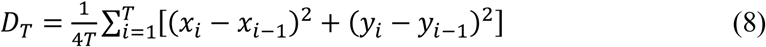

where *T* is the duration of the trajectory and *x*_*i*_ and *y*_*i*_ are the Cartesian position coordinates of the trajectory after time *i*. The effective diffusion coefficient was then related to the trajectory friction factor (*f*) by the Einstein relation (Edward, 1970):

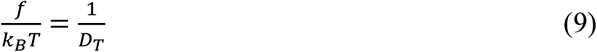

where *k*_*B*_ is the Boltzmann constant, *T* is temperature, and *D*_*T*_ is the effective diffusion coefficient for a single trajectory. For a more detailed explanation of experimental methods or trajectory friction factor calculations, see Thompson, et al (Thompson et al., 2019).

Considering that E-cad monomers tethered to a lipid diffuse the same as a single lipid at low surface coverage, we can extract the effective size of E-cad clusters assuming additive friction factor contributions from each E-cad molecule in the cluster (Cai et al., 2016; Knight et al., 2010; Thompson et al., 2019). The apparent trajectory friction factor, *f*, can be expanded as:

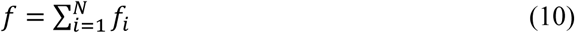

where *f*_*i*_ is the friction factor contribution due to each E-cad molecule in the cluster and N is the number of protein molecules in the cluster. The friction factor contribution of each protein is equal to the friction factor for an individual lipid in the free-draining limit, assuming bound lipids are well separated and each E-cad molecule tightly binds a single lipid and has minimal contact with additional lipids, all of which are true for this system after filtering trajectories (Knight et al., 2010). The lipid separation distance for this system should be nearly equal to the diameter of an E-cad extracellular domain, which is approximately 3.4 nm (Lambert et al., 2005; Nagar et al., 1996). This separation distance is large enough to assume lipid motion is not correlated (Knight et al., 2010). Therefore, the trajectory friction factor becomes:\

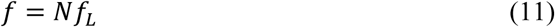

Where *f*_*L*_ is the friction factor of an individual lipid molecule diffusing in the supported lipid bilayer. The friction factor of an individual lipid molecule can be extracted from an E-cad friction factor distribution at low surface coverage by recognizing that the large peak in the low friction factor limit corresponds to E-cad monomer diffusion, and therefore, individual lipid diffusion. It was determined that *f*_*L*_ = 0.5 s/µm^2^ corresponded to the friction factor of a single lipid. The apparent cluster size was calculated for each trajectory, and a probability distribution was constructed for each experimental condition.

### Kinetic Monte Carlo Simulations

#### Construction of a domain-based coarse-grained model

Considering the E-cad extracellular regions consisting of five domains (EC1-EC5) (Oliver J. Harrison et al., 2011), we constructed a domain-based coarse-grained model to describe the structural arrangement of E-cad proteins. Each E-cad extracellular domain is coarse-grained into a rigid body with a radius of 1.5 nm, and the rigid bodies are spatially aligned into a rod-like shape (Fig. 4A). These E-cad extracellular domains are further distributed on the plasma membrane, which is represented by the bottom surface of a three-dimensional simulation box. The space above the plasma membrane represents the extracellular region. The extracellular regions of E-cad can form clusters through cis-interactions. Two different types of cis-interactions are considered in the model. The first is the polarized interactions that were observed in the crystal structure. To implement this interaction, we assigned a cis-donor site (purple dots) on the surface of each E-cad N-terminal domain, so that it can bind to a cis-receptor (red dots) site on the other E-cad. As a result, two adjacent E-cad proteins can be laterally connected through these specific cis-binding interfaces (Fig. 4A). In addition to the polarized specific interaction, a nonspecific interaction between two E-cads was also considered in the simulation system. As shown in the figure, this interaction can be formed by any pair of two E-cad within a certain distance cutoff. Therefore, it is non-polarized.

### Implementation of the kinetic Monte-Carlo (kMC) simulation algorithm

Given the surface density of E-cad, an initial configuration is constructed by randomly distributing molecules on the plasma membrane, as shown in Fig. 4B. Starting from this initial configuration, simulation of the dynamic system is then guided by a kinetic Monte-Carlo algorithm. The algorithm follows a standard diffusion-reaction protocol, as we developed earlier (Z.-R. Xie, J. Chen, & Y. Wu, 2014). Within each simulation time step, stochastic diffusions are first selected for randomly selected E-cad molecules. Translational and rotational movements of the molecules are confined on the surface at the bottom of the simulation box. The amplitude of these movements within each simulation step is determined by the diffusion coefficients of E-cad on a membrane surface. Periodic boundary conditions are implemented such that any E-cad that passes through one side of the cell surface reappears on the opposite side.

In conjunction with diffusion, the reaction associated with nonspecific and specific interactions is triggered stochastically if the binding criteria are satisfied between two E-cad molecules. The specific cis-interactions are triggered by two criteria: 1) the distance between a cis-donor site and a cis-receptor site of two molecules is below 1.2 nm cutoff (bond length), and 2) the orientation angles between two monomers are less than 30°, relative to the original configuration of the native E-cad dimer. Nonspecific interactions are trigged by one criterion: the distance between the center of mass of EC1 domains of two E-cad molecules is below 3.2 nm cutoff.

The probability of association is directly calculated by multiplying the on rate of the reaction with the length of the simulation time step. At the same time, dissociations are triggered for any randomly selected interaction with the probability that is calculated by multiplying the off rate of the corresponding reaction with the length of the simulation time step. If an E-cad molecule or E-cad cluster binds to another E-cad, or E-cad cluster through specific or nonspecific binding, they connect and move together subsequently on the surface of the plasma membrane. Finally, the above procedure is iterated until the system evolves into equilibrium patterns in both configurational and compositional spaces.

### Parameter determination in the coarse-grained simulations

The basic simulation parameters, including time step and binding criteria, were adopted from our previous work (Wang et al., 2018). The values of these parameters were determined based on benchmark tests in order to optimize the balance between simulation accuracy and computational efficiency. The two-dimensional translational diffusion constant of a single E-cad protein on a lipid bilayer is taken as 10 μm^2^/s and the rotational coefficient as 1° per ns. The values of these parameters were derived from our previous all-atom molecular dynamic simulation results for the diffusions of a cell-surface protein on the lipid bilayer (Z. R. Xie, J. Chen, & Y. Wu, 2014).

The reaction parameters, including the on and off rates of binding, were chosen from the range that is typical for protein-protein interactions, but at the same time make the simulations computationally accessible. As shown in the next section, the on rates for nonspecific and specific interactions are chosen from the range 10^8^ s^−1^ and 10^4^ s^−1^, corresponding to effective rate constants ranging from 10^4^ M^−1^s^−1^ to 10^8^ M^−1^s^−1^. This is a typical range for diffusion-limited rate constants, in which association is guided by complementary electrostatic surfaces at binding interfaces (Zhou & Bates, 2013). A wide range of off rates, from 10^4^ s^−1^ and 10 s^−1^, are used to model dissociation of both specific and nonspecific cis-interactions. Therefore, our tests cover the wide range of dissociation constants from milliMolar (mM) to nanoMolar (nM), which is within the typical range for binding of cadherin or other membrane receptors on cell surfaces.

### Sensitivity analysis

To evaluate the sensitivity of different parameters on E-cad clustering, we first performed kMC simulations at different E-cad concentrations. In order to exclude other factors, the on rate and off rates were fixed for nonspecific interactions at 2×10^−5^ s^−1^ and 10^3^ s^−1^, respectively, for both mutant and wild-type systems, and the on rate and off rate for specific interactions were fixed at 10^8^ s^−1^ and 10^2^ s^−1^, respectively, for wild-type systems. To build up the initial structure, we assign positions and orientations to 50, 100, 200 E-cad molecules on the membrane surface. The length of each side of the square plasma membrane surface is 400 nm, along both X and Y directions, which gives a total area of 0.16 µm^2^, leading to surface densities of 313 E-cad/µm^2^, 625 E-cad/µm^2^, 1,250 E-cad/µm^2^, respectively. At each concentration, we employed 50 independent replica simulations with random initial seeds. The simulations were extended to 0.8-1.3 seconds until the average cluster size reached equilibrium, and the final frames of trajectories were used for cluster size analysis. Fig. S7 shows resulting cluster size distributions at different concentrations. The solid lines represent one-term exponential fitting for each concentration. For comparison between experimental and simulated characteristic cluster size values, fitting was performed after removal of the data point corresponding to the bin at the smallest cluster size. The positive values of fitted characteristic cluster size (negative slope on semi-logarithmic plot), suggest that small cluster sizes are more favorable than large cluster sizes across the concentration range of 313 E-cad/µm^2^ to 1,250 E-cad/µm^2^. Meanwhile, our results show that large cluster sizes become more populated at higher E-cad concentration for both mutant and wild-type E-cad. This is consistent with experimental results showing that the characteristic cluster size increases with elevating E-cad concentration. Specifically, the characteristic cluster size for the cluster size distribution at 1,250 E-cad/µm^2^ is ∼29 E-cad, which is nearly the same for the experimental distribution at a concentration of 1,280 E-cad/µm^2^ (∼29 E-cad). These results indicate the robustness of our kMC simulation, suggesting that the clustering configuration generated from the model is sensitive to the total surface coverage of E-cad.

In order to further explore the sensitivity of the model to different binding parameters, we performed smaller kMC simulations involving various on- and off-rates of binding. To fix the surface density at 1,250 E-cad/µm^2^ and accelerate computing speed, we assigned only 50 E-cad molecules on a 100 × 100 nm^2^ membrane surface. For nonspecific interactions in mutant systems, the on rate values tested were 2×10^6^ s^−1^, 2×10^5^ s^−1^, and 2×10^4^ s^−1^, while the off rate values tested were 10^4^ s^−1^, 10^3^ s^−1^, and 10^2^ s^−1^. For specific interactions in wild-type systems, the on rate values tested were 10^8^ s^−1^, 10^7^ s^−1^, and 10^6^ s^−1^, and the off rate values tested were 10^3^ s^−1^, 10^2^ s^−1^, and 10 s^−1^, respectively. Simulations were carried out for all different combinations of on/off rates in the mutant system. At each on/off rate, we employed 10 or 20 independent runs with random initial seeds. The simulations were extended to 2 to 4 seconds, and the final 1s trajectories were used for cluster size analysis. Fig. S8 shows the effects of on/off rate on mutant E-cad cluster size distributions. In each panel, the solid red line represents a single exponential fit, and the values of the characteristic cluster sizes are shown in red. The panels with different on/off ratios have distinct characteristic cluster sizes, while the panels with the same on/off ratio (same binding affinity) have approximately the same characteristic cluster sizes. By comparing simulated and experimental characteristic cluster size values for the mutant, appropriate candidates of nonspecific on/off rates were identified. The optimum nonspecific on and off rates were 2×10^5^ s^−1^ and 10^3^ s^−1^, respectively. Using these nonspecific on/off rates, the wild-type system was simulated using all combinations of specific interaction on/off rates described above. Similarly, Fig. S9 shows the effects of specific interaction on/off rates on wild-type E-cad cluster size distributions. In each panel, the solid red line represents the single exponential fit, and the characteristic cluster size value is shown in red. The panels with different on/off rate ratios have distinct characteristic cluster sizes. Finally, analysis of simulated association time distributions can be utilized to select the best candidate from the combinations of specific on/off rates with the same ratio by comparing association time distributions for simulated and experimental trajectories. The selected on/off rates for nonspecific and specific interactions were the only combination of rates that resulted in qualitative agreement between simulated and experimental association time distributions.

## Additional Files

Supplementary File 1: Surface Residence Time Shows Interaction Dependence (Fig. S1).

Supplementary File 2: Additional Tables and Figures (Table S1 – Table S8 and Fig. S2 – Fig. S9).

## Acknowledgements

This work was supported by the National Institute of General Medical Sciences of the National Institutes of Health under award number 1R01GM117104.

---

**Supplementary File 1: Surface Residence Time Shows Interaction Dependence**

**Fig. S1.**
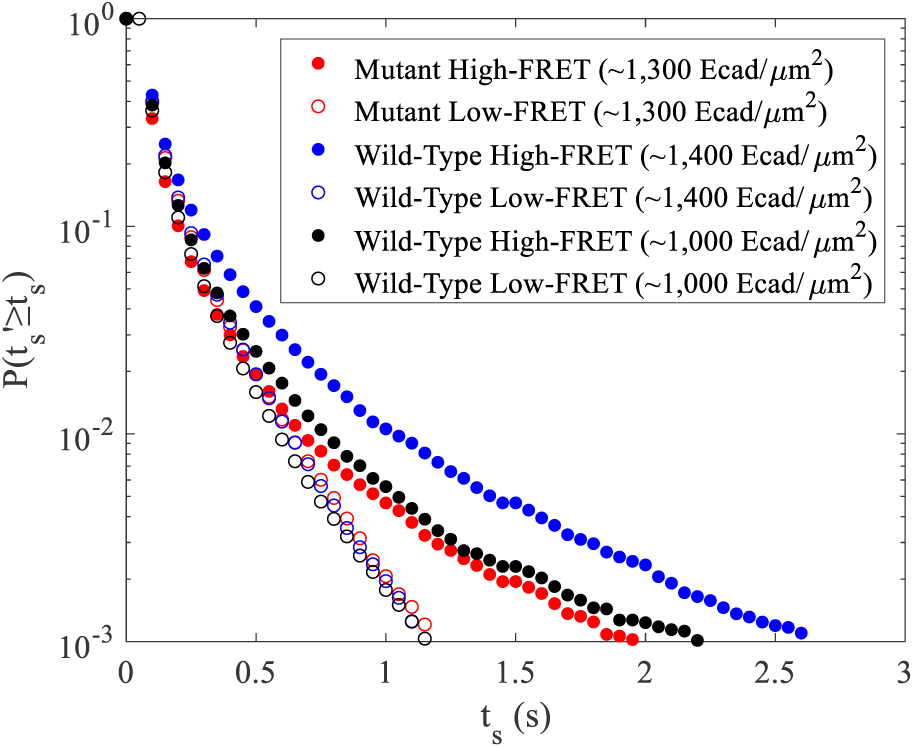
High-FRET and low-FRET complementary cumulative surface residence time distributions for mutant E-cad and two concentrations of wild-type E-cad.

Fig. S1 shows complementary cumulative surface residence time distributions for both low-FRET and high-FRET trajectories for the three experimental conditions. For additional details on complementary cumulative surface residence time distribution calculation in each FRET state, see the Methods section and the Supporting Information. Inspecting the low-FRET complementary cumulative surface residence time distributions for the mutant and two wild-type conditions, it is apparent that these distributions are not significantly different between these conditions. This implies that the surface residence times for low-FRET trajectories do not exhibit significant surface coverage dependence, as the high surface coverage wild-type condition is indistinguishable from the intermediate surface coverage wild-type condition. Additionally, comparing the high-surface coverage wild-type and mutant conditions, the inclusion of specific cis-interaction capability also does not significantly change the low-FRET trajectory surface residence times. Therefore, for low-FRET trajectories, the surface residence time distributions are primarily limited by the his-tag-NTA binding affinity and photobleaching. This is not the case for high-FRET trajectories, as it is apparent that the probability of a high-FRET trajectory having a long surface residence time is much increased compared to a low-FRET trajectory. The high-FRET surface residence time distributions also do exhibit surface coverage dependence for the wild-type, as indicated by the shift towards longer surface residence times when comparing the distributions for the high surface coverage wild-type and intermediate surface coverage wild-type condition. Furthermore, as a similar increase in surface residence time is seen when comparing the high surface coverage wild-type condition to the mutant condition, it is apparent that addition of cis-interactions causes an increase in surface residence times at comparable surface coverages. Overall, inspection of the high and low-FRET trajectory surface residence time distributions suggests that the surface residence time of a high-FRET trajectory was primarily limited by the protein interactions experienced by a molecule, instead of interaction lifetimes being limited by surface residence time. If this were not the case, one would expect that the high-FRET surface residence time distributions would be indistinguishable from the low-FRET distributions. This behavior allows a qualitative interpretation of the high-FRET state dwell time distributions, even though the dwell times are not necessarily bounded by transition to the low-FRET state.

**Supplementary File 2: Additional Tables and Figures**

**Table S1.** Number of trajectories and molecular observations used for diffusion analysis.

| Experimental Condition | Total Trajectories | Trajectories Exhibiting FRET | Total Molecular Observations |
| --- | --- | --- | --- |
| Wild-Type (~1,400 E-cad/$\mu\text{m}^2$) | 4,603 | 961 | 103,595 |
| Wild-Type (~1,000 E-cad/$\mu\text{m}^2$) | 4,628 | 799 | 102,196 |
| Mutant (~1,300 E-cad/$\mu\text{m}^2$) | 2,157 | 523 | 49,693 |

**Fig. S2.**
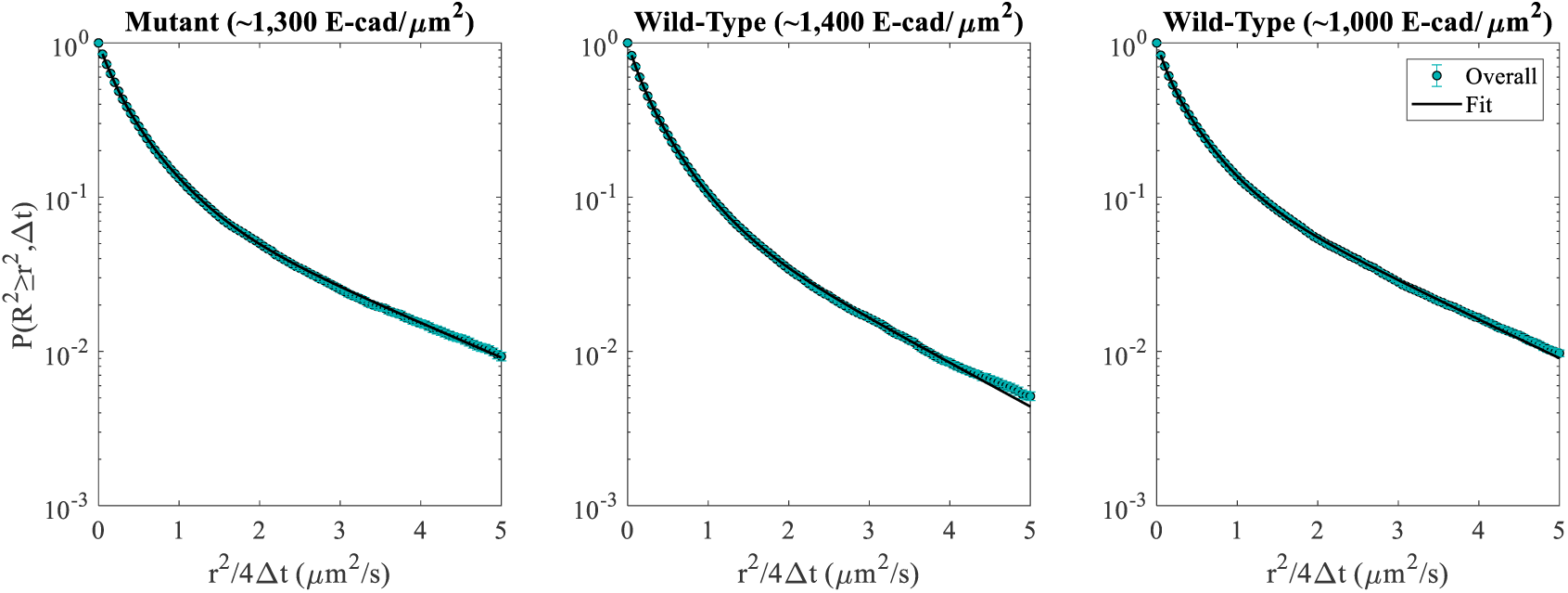
Overall CCSDDs, using all displacements from both high and low-FRET states, for the mutant and two wild-type E-cad conditions. Gaussian mixture model fits are shown as the black curve. Error bars represent the standard deviation of 100 bootstrapped samples.

**Table S2.** All CCSDD Gaussian mixture model fitting parameters for the overall, low-FRET, and high-FRET CCSDDs for the one mutant and two wild-type E-cad experimental conditions. Numbers in the parentheses represent the uncertainties (standard deviation of 100 bootstrapped samples) in the least significant digit.

| Experimental Condition | CCSDD | C<sub>1</sub> | C<sub>2</sub> | C<sub>3</sub> | D<sub>1</sub><br>($\mu\text{m}^2/\text{s}$) | D<sub>2</sub><br>($\mu\text{m}^2/\text{s}$) | D<sub>3</sub><br>($\mu\text{m}^2/\text{s}$) | $\overline{\text{D}}_{\text{short}}$<br>($\mu\text{m}^2/\text{s}$) |
| --- | --- | --- | --- | --- | --- | --- | --- | --- |
| Wild-Type<br>(~1,400 E-cad/$\mu\text{m}^2$) | Overall | 0.30(4) | 0.58(3) | 0.11(1) | 0.13(1) | 0.39(2) | 1.53(8) | 0.444(3) |
|  | Low-FRET | 0.36(7) | 0.50(6) | 0.13(2) | 0.17(2) | 0.43(4) | 1.7(1) | 0.494(5) |
|  | High-FRET | 0.31(4) | 0.59(3) | 0.10(3) | 0.10(1) | 0.38(3) | 1.2(2) | 0.365(4) |
| Wild-Type<br>(~1,000 E-cad/$\mu\text{m}^2$) | Overall | 0.28(3) | 0.56(2) | 0.16(1) | 0.119(7) | 0.40(2) | 1.73(6) | 0.537(5) |
|  | Low-FRET | 0.27(3) | 0.57(2) | 0.17(1) | 0.129(8) | 0.40(2) | 1.76(6) | 0.556(5) |
|  | High-FRET | 0.40(5) | 0.48(3) | 0.12(3) | 0.10(1) | 0.43(5) | 1.7(2) | 0.44(1) |
| Mutant<br>(~1,300 E-cad/$\mu\text{m}^2$) | Overall | 0.34(6) | 0.54(4) | 0.12(1) | 0.16(1) | 0.46(4) | 1.9(1) | 0.531(7) |
|  | Low-FRET | 0.40(5) | 0.50(4) | 0.10(2) | 0.19(1) | 0.54(4) | 2.3(2) | 0.569(8) |
|  | High-FRET | 0.15(5) | 0.70(4) | 0.14(1) | 0.07(2) | 0.31(2) | 1.5(1) | 0.44(1) |

**Fig. S3.**
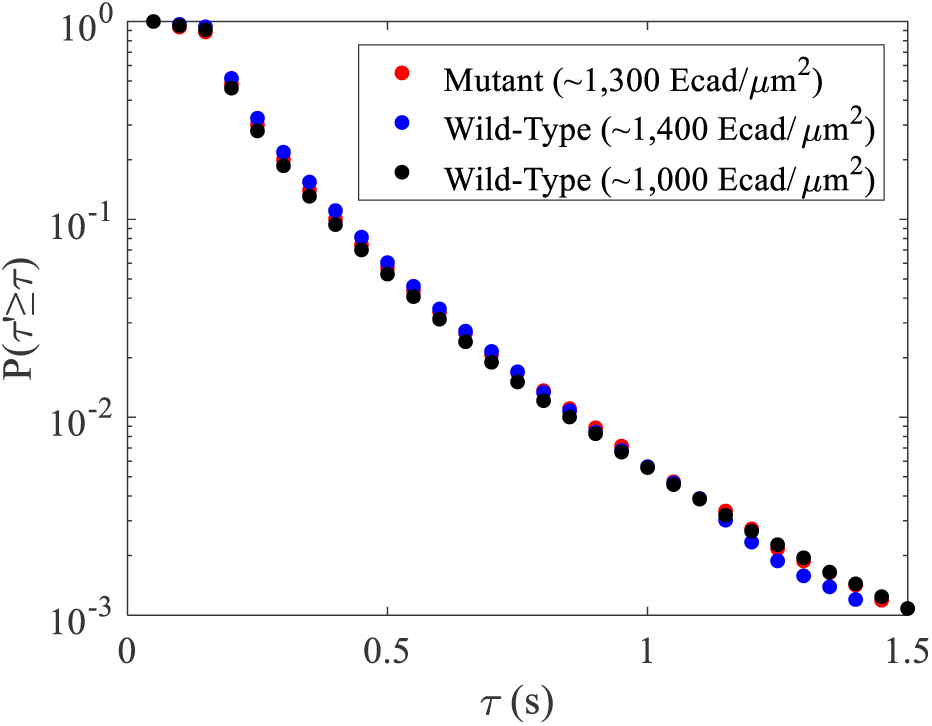
Low-FRET state complementary cumulative dwell time distributions for the mutant and two wild-type conditions.

**Fig. S4.**
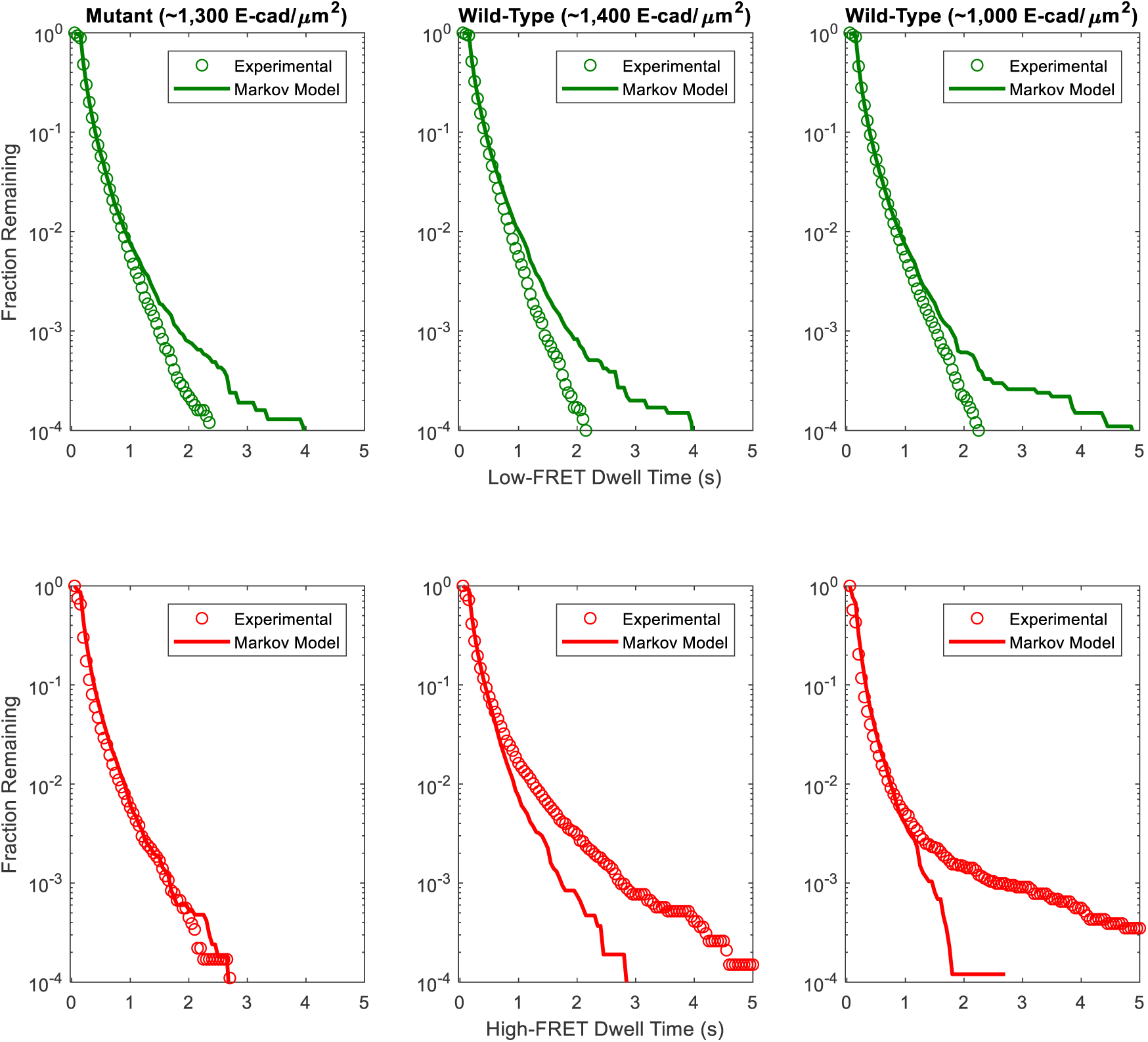
Complementary cumulative state dwell time distributions for the high and low-FRET states for the mutant and two wild-type E-cad conditions, compared to the predicted state dwell time distributions based upon the three-state, heterogeneous Markov model maximum likelihood estimate with beta-distributed transition probabilities.

**Table S3.**
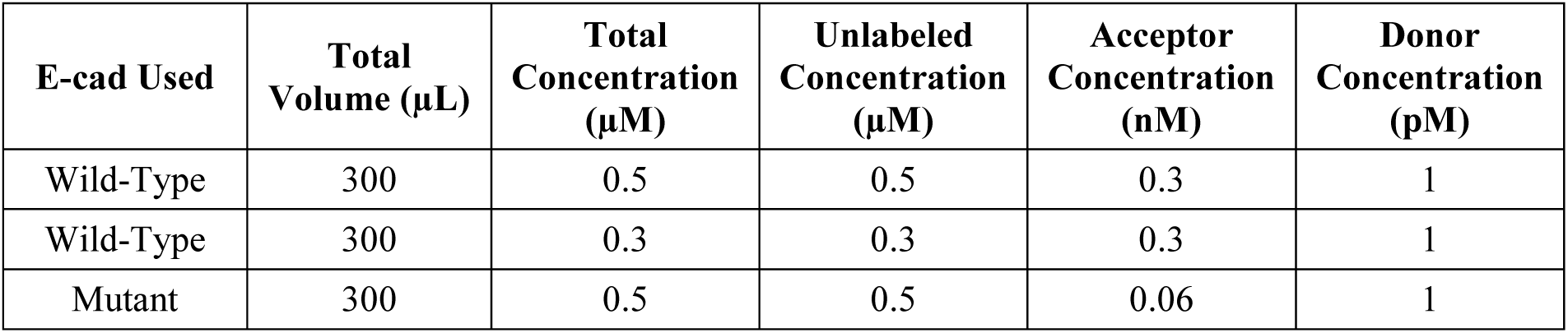
Protein solution concentrations for the two wild-type and one mutant E-cad conditions.

**Table S4.** Calculated two-dimensional surface coverage values for the two wild-type conditions with high and intermediate protein concentrations and the one mutant condition with high protein concentration. Numbers in parentheses represent the uncertainties (standard error between 12 or 13 different movies) in the least significant digit.

| Experimental Condition | Total Protein Solution Concentration ($\mu\text{M}$) | Surface Coverage (E-cad/$\mu\text{m}^2$) | Fractional Surface Coverage |
| --- | --- | --- | --- |
| Wild-Type | 0.5 | $1.37 \times 10^3(1)$ | 0.0124(1) |
| Wild-Type | 0.3 | $1.05 \times 10^3(2)$ | 0.0094(2) |
| Mutant | 0.5 | $1.26 \times 10^3(3)$ | 0.0113(2) |

**Fig. S5.**
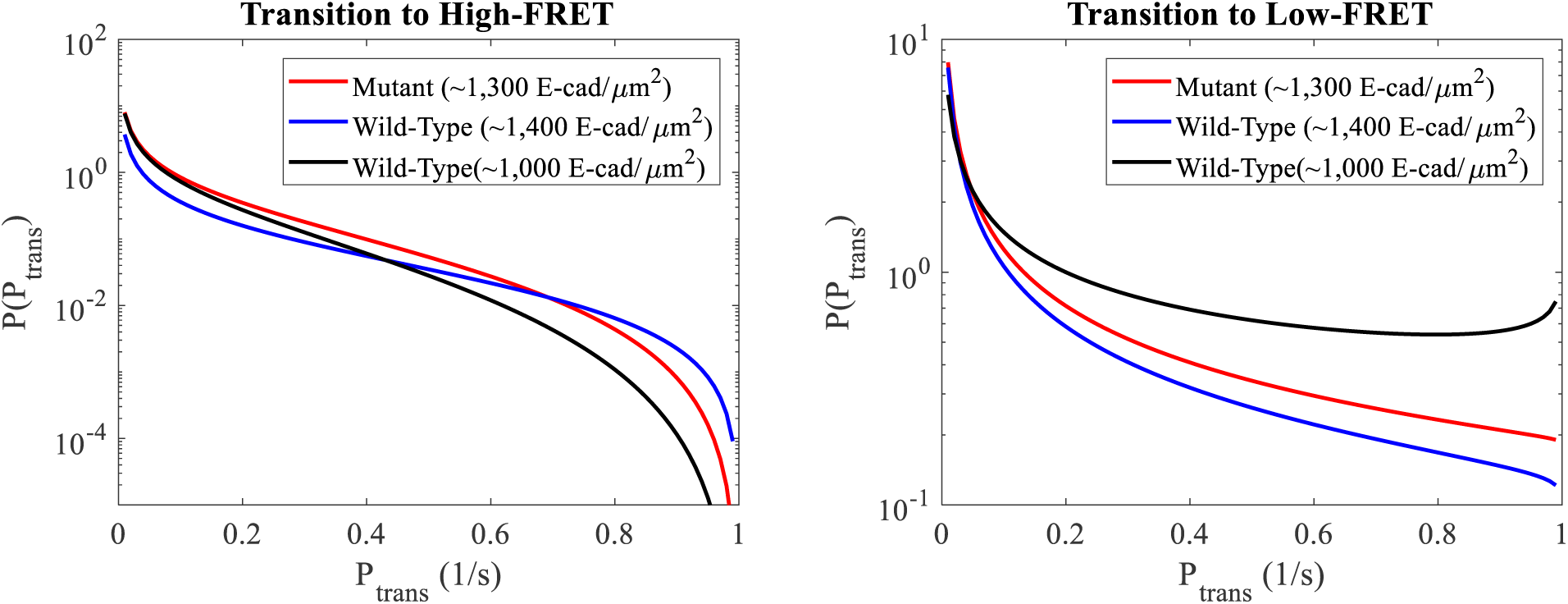
Beta distributions of state transition probabilities between the high and low-FRET states for the mutant and two wild-type conditions corresponding to the Markov model maximum likelihood estimated beta distribution parameters.

**Fig. S6.**
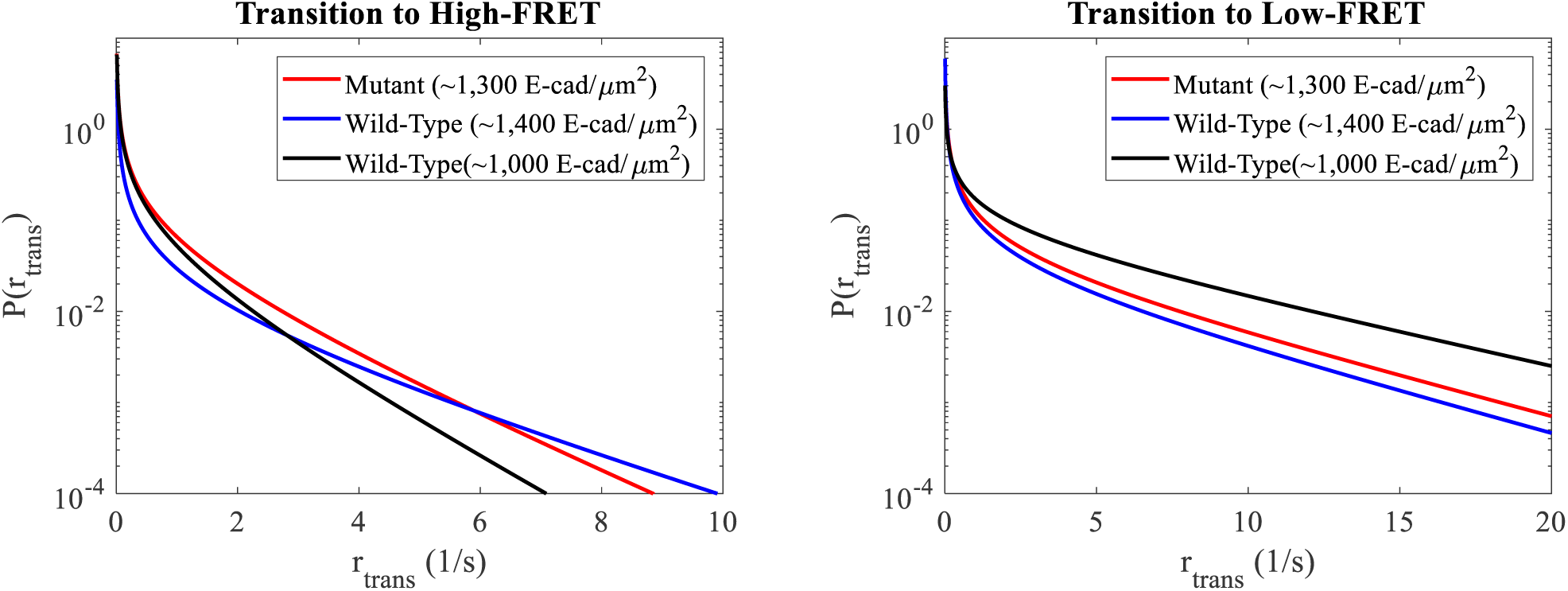
Probability density functions for the state transition rates between the high and low-FRET states for the mutant and wild-type conditions determined based upon the Markov model estimated, beta-distributed transition probabilities.

**Table S5.** Average transition rates between FRET states based on Markov model maximum likelihood estimated, beta-distributed transition probabilities. Numbers in parentheses represent the estimated uncertainties (square root of the Cramèr-Rao lower bound) in the least significant digit.

| Experimental Condition | Average Transition Rate<br>To High-FRET State<br>(1/s) | Average Transition Rate<br>To Low-FRET State (1/s) |
| --- | --- | --- |
| Wild-Type (~1,400 E-cad/$\mu\text{m}^2$) | 0.114(3) | 1.04(3) |
| Wild-Type (~1,000 E-cad/$\mu\text{m}^2$) | 0.145(3) | 3.17(6) |
| Mutant (~1,300 E-cad/$\mu\text{m}^2$) | 0.205(5) | 1.40(4) |

**Table S6.** System details for concentration sensitivity analysis.

| E-cad Used | Number of Proteins | Surface Area (nm<sup>2</sup>) | Concentration (E-cad/$\mu\text{m}^2$) | Number of Simulation Runs | Average Cluster Size (# of E-cad) |
| --- | --- | --- | --- | --- | --- |
| Wild-Type | 200 | 400 × 400 | 1,250 | 50 | 39.6 |
| Wild-Type | 100 | 400 × 400 | 625 | 50 | 26.0 |
| Wild-Type | 50 | 400 × 400 | 313 | 50 | 19.6 |
| Mutant | 200 | 400 × 400 | 1,250 | 50 | 28.4 |
| Mutant | 100 | 400 × 400 | 625 | 50 | 19.0 |
| Mutant | 50 | 400 × 400 | 313 | 50 | 9.0 |

**Table S7.** System details for on/off rate sensitivity analysis.

| E-cad Used | Number of Proteins | Concentration (E-cad/$\mu\text{m}^2$) | Nonspecific Interaction On/off Rate (s<sup>-1</sup>) | Specific Interaction On/off Rate (s<sup>-1</sup>) | Number of Simulation Runs |
| --- | --- | --- | --- | --- | --- |
| Mutant | 50 | 1,250 | $2 \times 10^6 / 1 \times 10^4$ | N/A | 10 |
| Mutant | 50 | 1,250 | $2 \times 10^6 / 1 \times 10^3$ | N/A | 10 |
| Mutant | 50 | 1,250 | $2 \times 10^6 / 1 \times 10^2$ | N/A | 10 |
| Mutant | 50 | 1,250 | $2 \times 10^5 / 1 \times 10^4$ | N/A | 10 |
| Mutant | 50 | 1,250 | $2 \times 10^5 / 1 \times 10^3$ | N/A | 10 |
| Mutant | 50 | 1,250 | $2 \times 10^5 / 1 \times 10^2$ | N/A | 10 |
| Mutant | 50 | 1,250 | $2 \times 10^4 / 1 \times 10^3$ | N/A | 10 |
| Mutant | 50 | 1,250 | $2 \times 10^4 / 1 \times 10^2$ | N/A | 10 |
| Wild-Type | 50 | 1,250 | $2 \times 10^5 / 1 \times 10^3$ | $1 \times 10^8 / 1 \times 10^3$ | 20 |
| Wild-Type | 50 | 1,250 | $2 \times 10^5 / 1 \times 10^3$ | $1 \times 10^8 / 1 \times 10^2$ | 20 |
| Wild-Type | 50 | 1,250 | $2 \times 10^5 / 1 \times 10^3$ | $1 \times 10^8 / 1 \times 10$ | 20 |
| Wild-Type | 50 | 1,250 | $2 \times 10^5 / 1 \times 10^3$ | $1 \times 10^7 / 1 \times 10^3$ | 10 |
| Wild-Type | 50 | 1,250 | $2 \times 10^5 / 1 \times 10^3$ | $1 \times 10^7 / 1 \times 10^2$ | 10 |
| Wild-Type | 50 | 1,250 | $2 \times 10^5 / 1 \times 10^3$ | $1 \times 10^7 / 1 \times 10$ | 10 |
| Wild-Type | 50 | 1,250 | $2 \times 10^5 / 1 \times 10^3$ | $1 \times 10^6 / 1 \times 10^2$ | 10 |
| Wild-Type | 50 | 1,250 | $2 \times 10^5 / 1 \times 10^3$ | $1 \times 10^6 / 1 \times 10^2$ | 10 |
| Wild-Type | 50 | 1,250 | $2 \times 10^5 / 1 \times 10^3$ | $1 \times 10^6 / 1 \times 10$ | 10 |

**Fig. S7.**
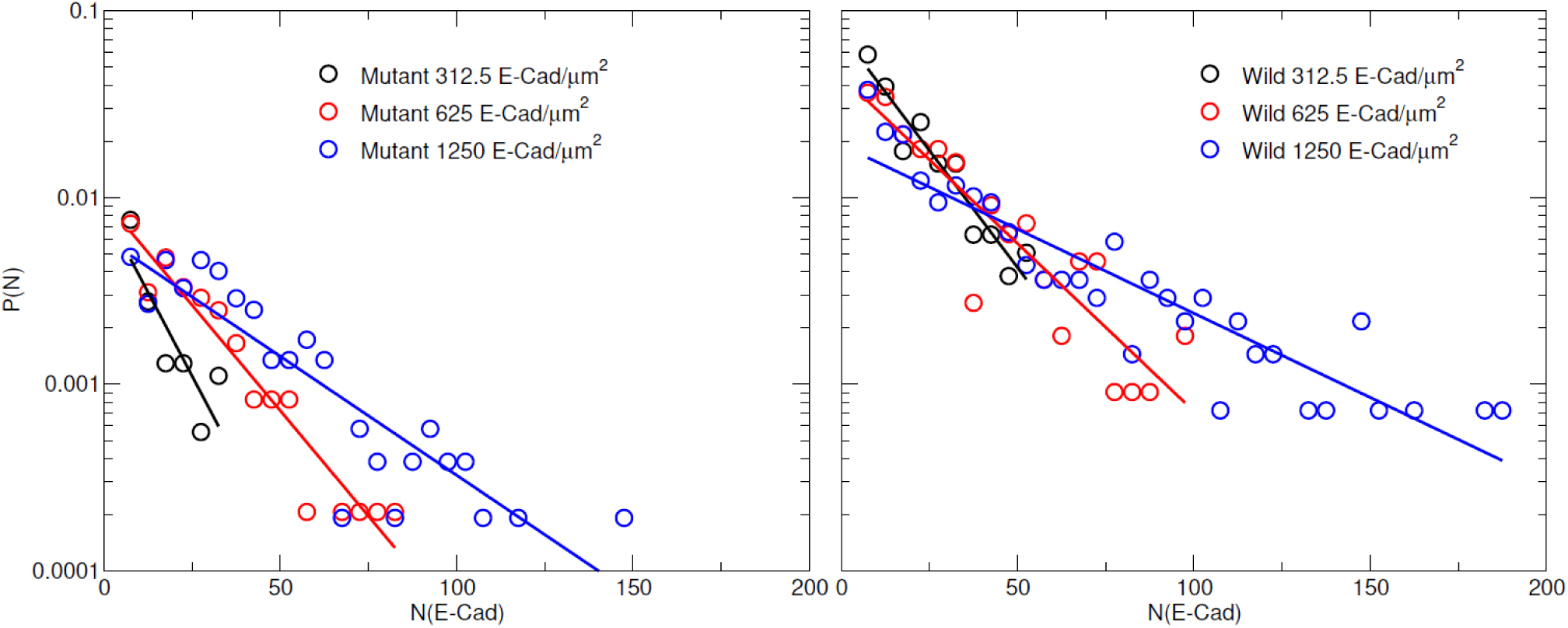
Distribution of cluster size for wild-type and mutant E-cad at a surface density of 312.5 E-cad/µm^2^, 625 E-cad/µm^2^, and 1,250 E-cad/µm^2^, respectively. The data are fit to a single exponential.

**Fig. S8.**
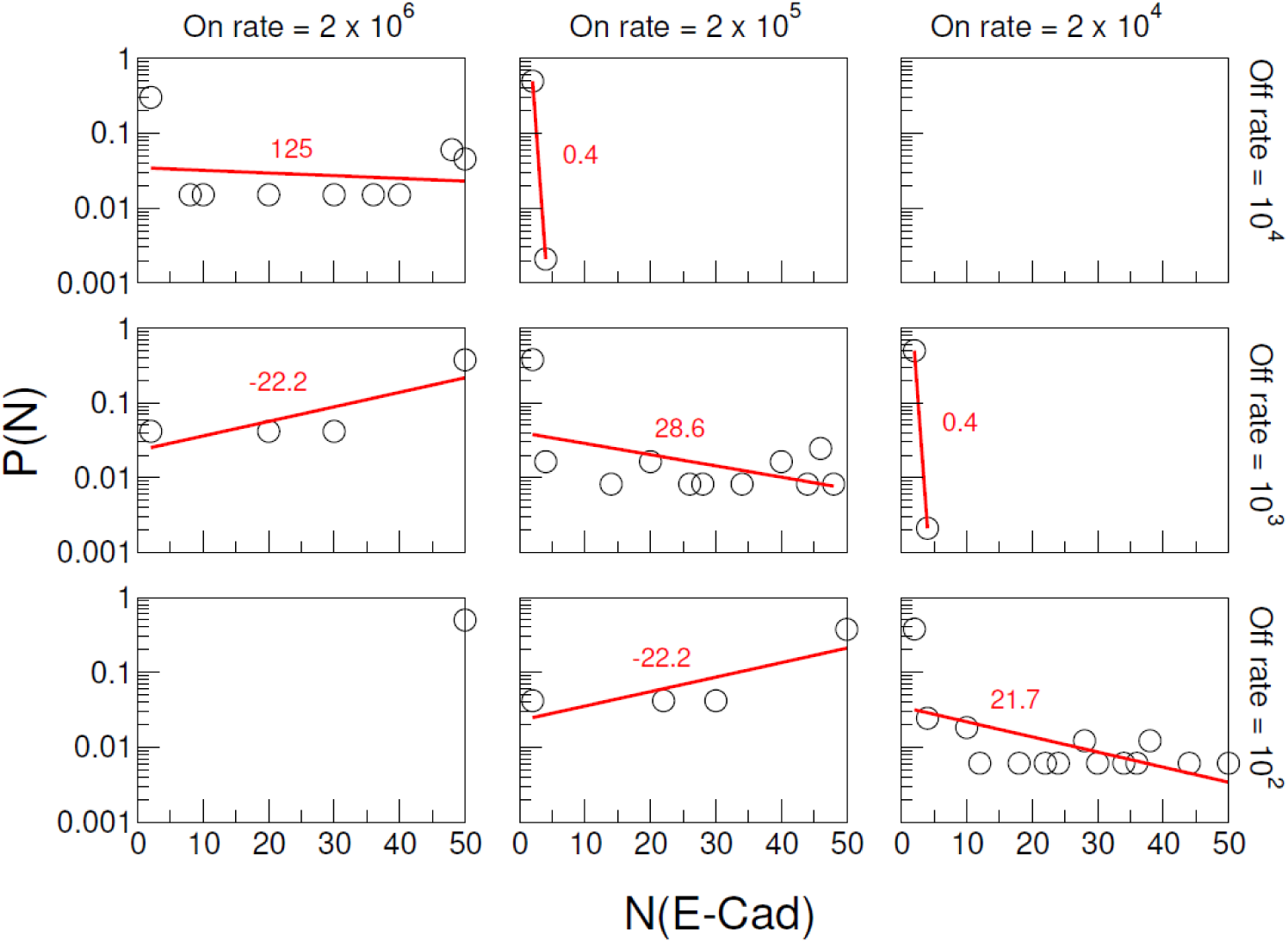
Distributions of mutant E-cad cluster size for different combinations of nonspecific interaction on/off rate. The 1^st^, 2^nd^ and 3^rd^ columns correspond to on rate 2×10^6^ s^−1^, 2×10^5^ s^−1^, and 2×10^4^ s^−1^, respectively. The 1^st^,2^nd^ and 3^rd^ rows correspond to off rate 10^4^ s^−1^, 10^3^ s^−1^, and 10^2^ s^−1^, respectively. In each panel, the cluster size distributions are fitted by a single exponential function 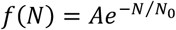 with a constant N_0_ corresponding to the characteristic cluster size. The fitted line and N_0_ values are colored in red.

**Fig. S9.**
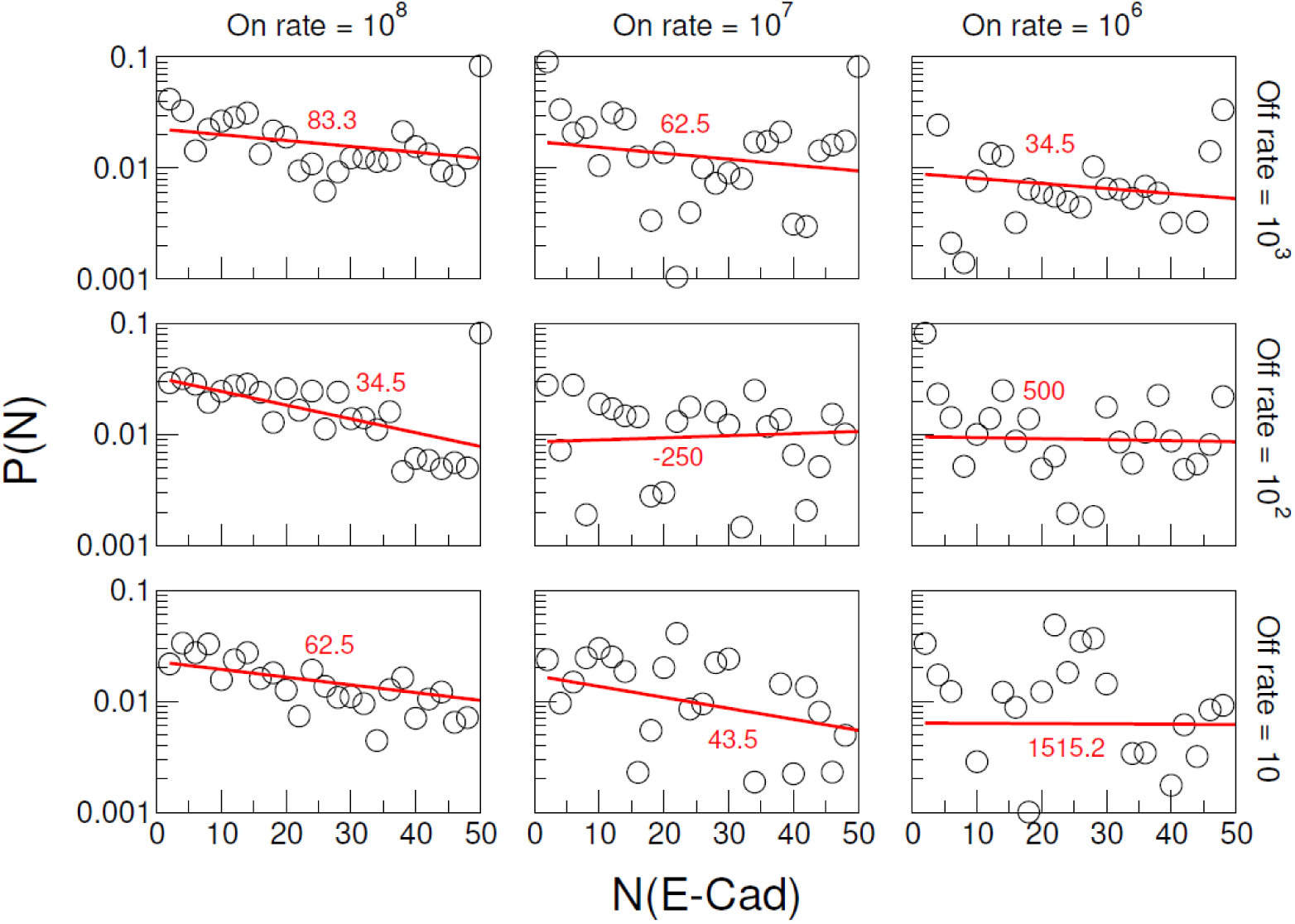
Distributions of wild-type E-cad cluster size for different combinations of specific interaction on/off rate. The columns correspond to on rate 10^8^ s^−1^, 10^7^ s^−1^, and 10^6^ s^−1^, respectively. The rows correspond to off rate 10^3^ s^−1^, 10^2^ s^−1^, and 10 s^−1^, respectively. Nonspecific interaction on and off rates, respectively, were fixed at 2×10^5^ s^−1^ and 10^3^ s^−1^. In each panel, the cluster size distributions are fitted by a single exponential function 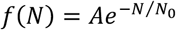 with a constant N_0_ corresponding to the characteristic cluster size. The fitted line and N_0_ value are colored in red.

**Table S8.** The rates used in kMC simulations that best fit the experimental cluster size distribution and experimental association time distributions.

| | On Rate ($\text{s}^{-1}$) | Off Rate ($\text{s}^{-1}$) |
| --- | --- | --- |
| Nonspecific interaction | $2 \times 10^5$ | $10^3$ |
| Specific interaction | $10^8$ | $10^2$ |

